# A novel PNGase Rc for improved protein N-deglycosylation in bioanalytics and HDX-MS epitope mapping under challenging conditions

**DOI:** 10.1101/2022.04.13.488165

**Authors:** Marius Gramlich, Sandra Maier, Philipp D. Kaiser, Bjoern Traenkle, Teresa R. Wagner, Josef Voglmeir, Dieter Stoll, Ulrich Rothbauer, Anne Zeck

## Abstract

N-linked glycosylation is a ubiquitous posttranslational modification of proteins. While it plays an important role in the biological function of proteins, it often poses a major challenge for their analytical characterization. Currently available peptide N-glycanases (PNGases) are often inefficient at deglycosylating proteins due to sterically inaccessible N-glycosylation sites. This usually leads to poor sequence coverage in bottom-up analysis using liquid chromatography with tandem mass spectrometry (LC-MS) and makes it impossible to obtain an intact mass signal in top-down MS analysis. In addition, most PNGases operate optimally only in the neutral to slightly acidic pH range and are severely compromised in the presence of reducing and denaturing substances, which limits their use for advanced bioanalysis based on hydrogen-deuterium exchange in combination with mass spectrometry (HDX-MS).

Here, we present a novel peptide N-glycanase from *Rudaea cellulosilytica* (PNGase Rc) for which we demonstrate broad substrate specificity for N-glycan hydrolysis from multiply occupied and natively folded proteins. Our results show that PNGase Rc is functional even under challenging, HDX quench conditions (pH 2.5, 0 °C) and in the presence of 0.4 M Tris(2-carboxyethyl)phosphine (TCEP), 4 M urea and 1 M guanidinium chloride (GdmCl). Most importantly, we successfully applied the PNGase Rc in an HDX-MS workflow to determine the epitope of a nanobody targeting the extracellular domain of human signal-regulating protein alpha (SIRPα).

## Introduction

Numerous proteins undergo post-translational modifications (PTMs), including disulfide bond formation and glycosylation, which confer proper folding, solubility, and stability to their native conformation in order to maintain correct biological function^1–4^. Protein glycosylation is involved in a variety of biological processes, such as cellular communication, receptor activation, tumor growth and metastasis, and viral evasion of the immune system^4–6^. Thus, extracellular glycosylated proteins represent attractive drug targets or are themselves used as biopharmaceuticals^3, 7, 8^. Protein glycosylation can be mainly divided into two groups, the O- and the N-linked glycans. O-linked oligosaccharides are attached to the hydroxyl oxygen of threonine or serine residues, whereas N-linked glycans are attached to the nitrogen of asparagine side chains through an N-glycosidic bond^2^. The percentage occupancy at a given site (macroheterogeneity) by different individual glycan endowments (microheterogeneity) generally results in very heterogeneous PTM profiles^7, 8^. This in turn leads to a distribution of the signal obtained, e.g. in mass spectrometry, over several molecular species, which impedes the analysis at protein and peptide level^9, 10^.

Hydrogen deuterium exchange coupled with mass spectrometry (HDX-MS) is a powerful approach that can provide insights into protein behavior by serving as a link between structure, conformational dynamics and function^11^. The technique became highly attractive for obtaining information on protein dynamics, protein-drug interactions, epitope characterization or for routine quality control within the biopharmaceutical industry^12–16^. As a chemical footprinting technique, it measures the exchange of amide hydrogen atoms in the protein backbone against heavier deuterium atoms from deuterated solvents. The exchange rate is mainly influenced by the solvent accessibility of the amide protons and their involvement in hydrogen bonds. HDX can be performed at the global protein or peptide level, using a bottom-up approach including a proteolysis step^17, 18^. Since HDX is a reversible process sensitive to temperature and pH, sample treatment must be performed within a short period of time and at a pH of 2.5 and low 0 °C (HDX quench conditions)^19^. Compared to other structural analysis methods, HDX-MS shows high sensitivity and tolerance towards sample complexity. However, PTMs such as disulfide bonds and N-glycosylation pose a particular challenge for HDX analysis because they impede peptide identification due to signal distribution across multiple glycopeptide species with unknown N-glycan content and non-specifically cleaved peptic peptide sequence^9, 20, 21^. Furthermore, mass spectrometry (MS) data derived from glycopeptides using conventional fragmentation techniques such as collision-induce dissociation (CID) are difficult to interprete because little information about the peptide backbone is obtained. In CID, the weak glycosidic bonds are preferentially fragmented compared to the amide bonds of the peptides^22^. Therefore, HDX-MS often shows a lack of sequence coverage and deuteration information in regions containing N-linked glycosylation sites^9, 23, 24^. If the peptide sequence can be identified anyway, glycosylation still represents a source of misinterpretation of HD exchange data because the N-glycan pentasaccharide core contains acetamido groups, which can exchange protons and retain deuterium even under quenching conditions^25^.

Enzymatic N-glycan release by peptide-N(4)-(N-acetyl-β-glucosaminyl)asparagine amidases (PNGases) is a valuable approach to lower the heterogeneity and subsequently facilitate the analysis of N-glycosylated proteins^7, 26–28^. Two commercially available PNGases derived either from *Flavobacterium meningosepticum* (PNGase F)^29^ or *Prunus dulcis* (PNGase A)^30^ are widely used for deglycosylation. Notably, PNGase F has a physiological pH working range and shows no activity below pH 4.0^29^, which limits its use for HDX-MS to deglycosylation prior to deuterium labeling. However, such a workflow could lead to a structural rearrangement in the investigated protein and thus prevents analysis of the protein in its native conformation^23, 31^. In contrast, PNGase A was successfully used under quench conditions for post-HDX deglycosylation due to its residual activity at pH 2.5^32, 33^.

In addition to glycosylation, disulfide bonds pose an additional challenge for HDX-MS because they interfere with proteolysis, resulting in a lack of sequence coverage near the cysteine residues involved in the disulfide bonds. Reduction of disulfide bonds is preferentially achieved with tris-(2-carboxyethyl)phosphine (TCEP), but this works inefficiently at the low temperatures, low pH values, and short reaction times required for HDX experiments. Therefore, additional steps to denature the proteins by adding chaotropic agents, such as urea or guanidine hydrochloride (GdmCl) are usually required^17, 34–36^ which severely compromises the functionality of the applied PNGase as been reported for acidic PNGase A^33^. More recently, a set of novel acidic PNGase enzymes has been discovered by Guo *et al*.^37^ and successfully applied in an online HDX workflow to study an extracellular protein comprising multiple disulfide bonds and N-glycosylated residues^38^. However, with the electrochemical reduction of disulfide bonds and a novel PNGase variant immobilized on microfluidic chips, this study required a complex experimental setup and is further limited by the poor availability of the enzyme used.

Here we present a novel acidic PNGase variant from *Rudaea cellulosilytica* (PNGase Rc). This variant can be efficiently produced and purified in high yields upon heterologous expression in *Escherichia coli* (*E.coli*) and reveals a high deglycosylation efficiency at pH 2.5 at intact protein level. For functional studies, we implemented the PNGase Rc into our semi-automatic HDX-MS bottom-up workflow, which enables N-glycan release of peptic peptides post-labeling under harsh acidic, denaturing and reducing conditions. Our findings show that the enzyme is highly resistant to high concentrations of TCEP and urea and shows tolerance against guanidine hydrochloride (GdmCl). In a proof-of-principle study, we successfully used PNGase Rc in an HDX-MS workflow and elucidated the epitope of a single domain antibody fragment (nanobody) raised against the multiple N-glycosylated extracellular domain of the human signal-regulatory protein alpha (SIRPα).

## Results

### High yield heterologous expression of PNGase Rc

A prerequisite for the efficient production of glycanases, which often originate from rare prokaryotes, is the transfer of the encoding cDNA into an established bacterial expression system. Therefore, we cloned the ORF encoding for PNGase Rc from the original soil bacterium *Rudaea cellulosilytica*^39^ in a bacterial expression vector (pET30a(+)), thereby adding an N-terminal His_6_-tag (**Figure S1**). Heterologous expression was carried out in *Escherichia coli* strain BL21 (DE3) and the recombinant enzyme was isolated and purified using immobilized metal ion affinity chromatography (IMAC) followed by size exclusion chromatography (SEC) yielding ∼1.3 mg of purified enzyme with a purity of ∼75% from 1L *E.coli* culture (**Figure 1a**). Subsequently, we examined the identity and integrity of the recombinant PNGase Rc by electrospray MS (**Figure 1b**). Deconvolution of the mass-to-charge signal series confirms the theoretical mass of 64198.0 Da with closed disulfide links. A minor side peak at 59129.7 Da corresponds to a truncated protein variant lacking the first 47 N-terminal amino acids (59128.5 Da), which confers to the common autohydrolysis amino acid motif “DP” and may originate from the sample itself or may be artificially introduced by electrospray ionization. The additional peaks at 20948.2 Da and 23508.3 Da could not be assigned to the amino acid sequence of the enzyme and might derived from contaminant proteins detected in the SDS-PAGE at similar molecular weight (**Figure 1a**).

**Figure 1.**
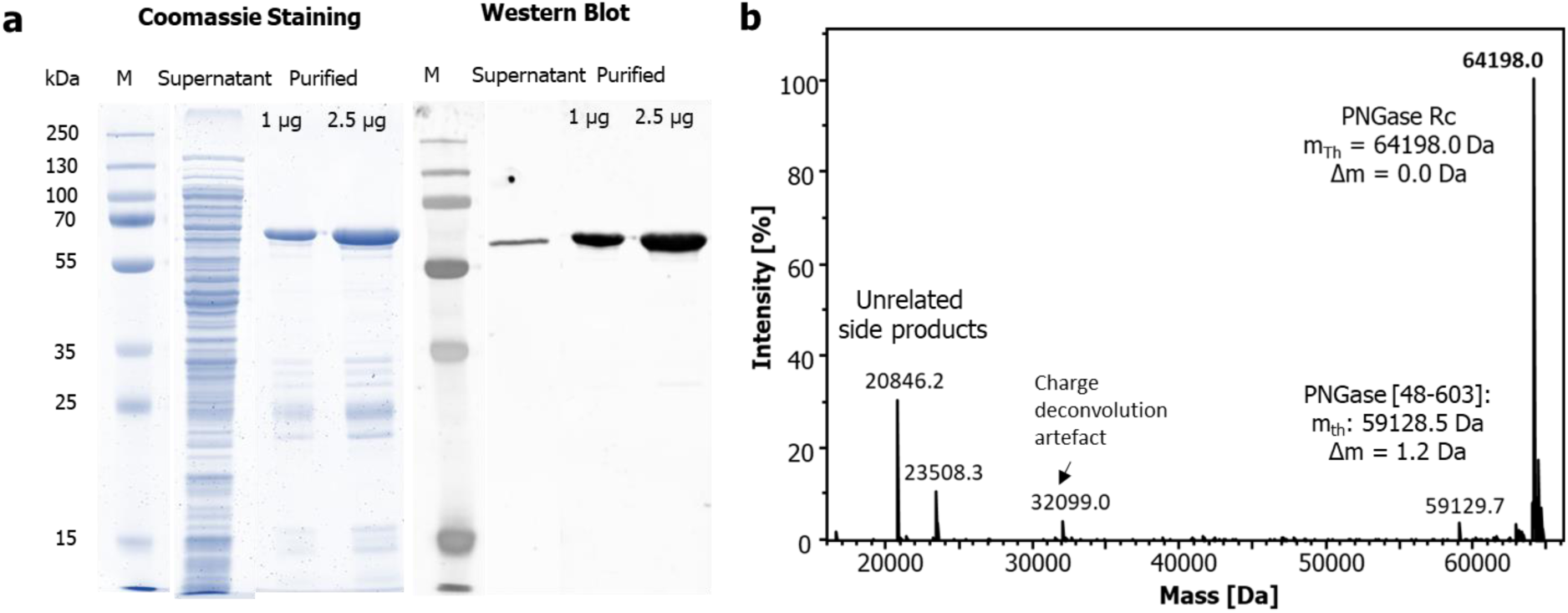
Purification and mass analysis of the recombinant PNGase from *Rudaea cellulosilytica* (Rc). (**a**) SDS-PAGE followed by coomassie staining (left) and western blot analysis (right) of the PNGase Rc purified from *E.coli*. Western blot analysis was performed using an anti-His-tag antibody in combination with a fluorescently-labeled anti-mouse antibody. (**b**) Identity and integrity confirmation of the recombinant PNGase Rc using intact mass spectrometry (MS) analysis.

### Determination of PNGase Rc enzyme activity

To evaluate the activity of recombinant PNGase Rc, we employed a mass spectrometric assay, monitoring the hydrolysis of N-glycan units of intact trastuzumab (human IgG1 antibody) upon incubation with PNGase Rc in a time-dependent manner. Each of the two heavy chains of the IgG molecule is fully occupied by an N-glycan at asparagine (N) 301 showing no macroheterogeneity. With respect to the microheterogeneity, N301 is occupied mainly by three glycoforms containing complex biantennary glycans with different degrees of galactosylation (**Figure 2a**)^40^. These glycoforms were summed up for assessment of the deglycosylation reaction. Deglycosylation was performed using a molar enzyme-to-substrate ratio (E:S) of 1:44 at pH 2.5 and 37 °C and abundances of the non-, half and fully deglycosylated antibody molecules were monitored and relatively quantified after different time periods for up to 40 min (**Figure 2a**). Only the degree of fully deglycosylated antibody was considered for the determination of the reaction kinetics. Our results showed 41% and 100% fully deglycosylated intact antibody after 10 and 40 min incubation, respectively (**Figure 2b**). Additionally, we compared the activity of the novel PNGase Rc with the commercially available PNGase F, which is widely used for deglycosylation of intact proteins^27^. For this, we performed the established assay and only changed the pH to 7.5, which is within the optimal working range of PNGase F (**Figure 2b**). Compared to PNGase Rc, a significantly lower N-glycan release rate was observed for PNGase F, resulting in a deglycosylation of only 15 % after 40 min of incubation. The determined reaction rate 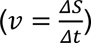 of the PNGase Rc is 5.2 %/min and therefore ∼10 times higher than that of PNGase F (0.4 %/min) (**Figure 2b**). This corresponds to a molar N-glycan hydrolysis rate of 17.7 and 1.4 pmol/min for PNGase Rc and F, respectively. From these findings, we concluded that PNGase Rc exhibits not only a high intrinsic activity but also efficiently removes complex biantennary N-glycans with core fucosylation.

**Figure 2.**
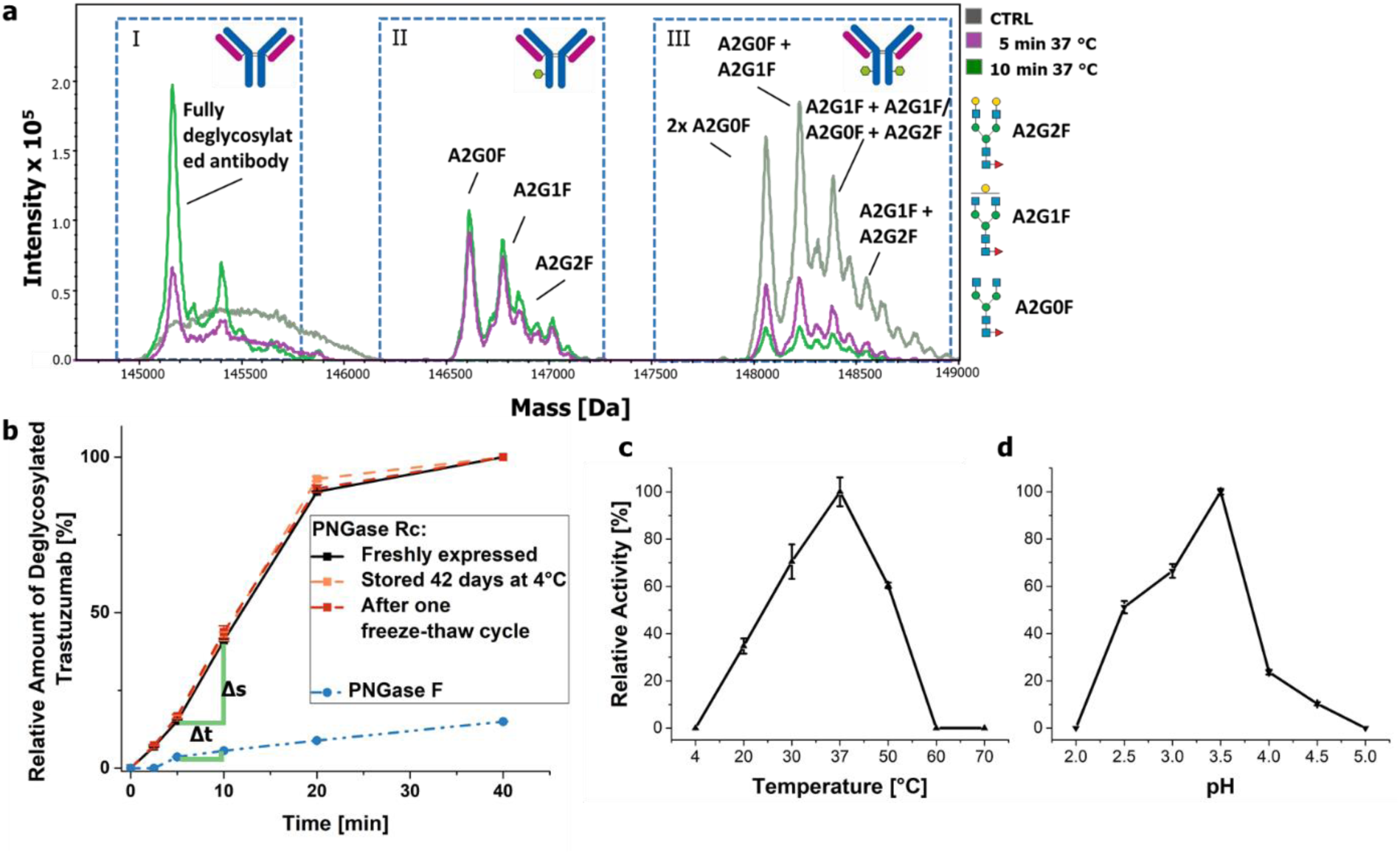
PNGase activity assay monitoring deglycosylation of an intact antibody by mass spectrometry (a, b), temperature (c) and pH working range (d). (**a**) Overlay of charge-deconvoluted mass spectra showing fully (I), half (II) and non-(III) deglycosylated IgG molecules after incubation with PNGase Rc. (**b**) Relative amount of fully deglycosylated IgG using PNGase Rc after different storage conditions and compared to PNGase F deglycosylation (T = 37 °C; E:S 1:44; pH 3.5 (PNGase Rc) and pH 7.5 (PNGase F)). (**c**) Full deglycosylation of intact IgG with PNGase Rc (t = 10 min, pH 2.5). (**d**) Full deglycosylation of intact IgG with PNGase Rc (t = 10 min; T = 37 °C; different pH). Shown is the ±SEM of three technical replicates.

### Storage stability, pH and temperature optimum of PNGase Rc

Next, we analyzed important parameters such as the enzyme storage stability under different conditions. Therefore, we examined the enzyme activity following extended storage at 4 °C for 42 days and after one freeze-thaw cycle using the described protein assay. According to our data no significant decrease in PNGase activity for any of the selected treatments was observable (**Figure 2b**). Furthermore, we determined temperature and pH working ranges of the enzyme in independent technical triplicates. Results from these experiments are summarized in **Figure 2c-d** with the observed activities normalized to the highest detected enzyme activity. Starting from initial conditions at pH 2.5 and 37 °C, both parameters were changed separately and incrementally using a fixed incubation time of 10 min and enzyme/substrate molar ratio of 1:44. Our measurements revealed a temperature optimum of 37 °C, while higher temperatures above 50 °C led to inactivation of the PNGase Rc (**Figure 2c**). Notably, we did not observe full deglycosylation in the natively folded antibody at 4 °C (on-ice) within a 10 minute time range under the assay conditions. However, 6.3% half deglycosylated antibody species indicated remaining enzyme activity at this low temperature. Analyzing different pH conditions further revealed an activity optimum at pH 3.5 with a working range of pH 2.5 - 4.5 and a complete inactivation of the PNGase Rc activity outside this range (**Figure 2d**).

### Substrate specificity of PNGase Rc for native protein N-deglycosylation compared to PNGase F

To further evaluate the substrate specificity of PNGase Rc, we examined the ability of the enzyme to release N-glycans for three multiply or specifically glycosylated proteins, namely ribonuclease B (RNase B), horseradish peroxidase (HRP), and fetuin in a time-dependent manner. As a reference, we used PNGase F and deglycosylation was monitored by the band shift obtained using SDS-PAGE (**Figure 3a I-III**). For all experiments we used identical E:S ratios and optimal pH conditions for each enzyme. Additionally we applied protein denaturation prior to deglycosylation, which is known to facilitate deglycosylation of sterically inaccessible sites^27^.

**Figure 3.**
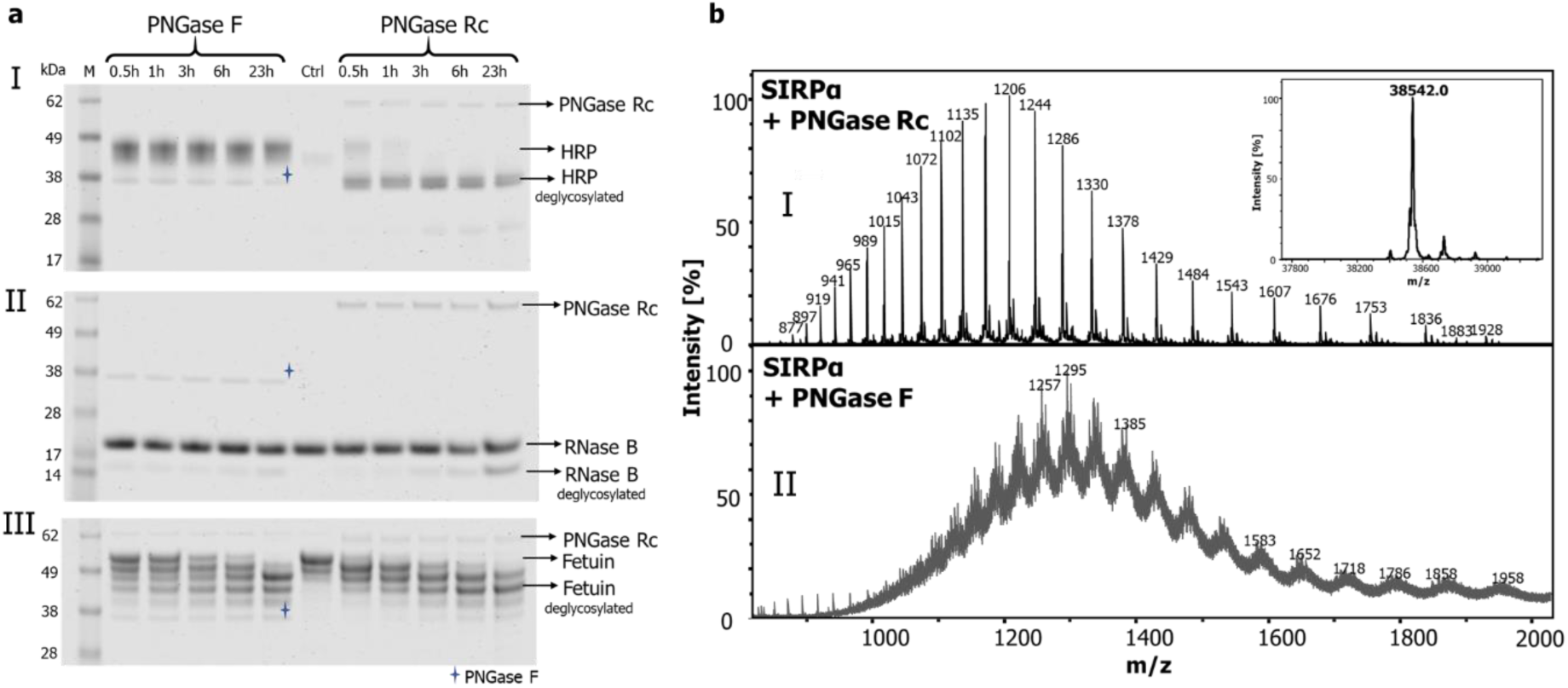
Comparison of intact protein deglycosylation by PNGase Rc and F tested with HRP, RNase B and fetuin and the multiply glycosylated extracellular receptor domain of human SIRPα. (**a**) HRP (I), RNase B (II) or fetuin (III) were incubated for various times with PNGase Rc or F at pH 3.5 or 7.5, respectively. Deglycosylation was performed at an E:S of 1:48 (M:M) and at 37 °C. Deglycosylation efficiencies were monitored by SDS-PAGE followed by coomassie staining at indicated time points. (**b**) m/z spectra of intact SIRPα after overnight incubation at 37 °C and an E:S of 1:45 with PNGase Rc (I) or PNGase F (II), at pH 2.5 or pH 7.4 respectively. The inlayer of the lower panel shows the charge-deconvoluted mass spectrum confirming the identity of SIRPα.

HRP (∼44 kDa) contains nine predicted N-glycosylation sites, which for the most part (> 80%)^41^ are occupied by typical plant N-glycans (MMXF) bearing α-1,3-core-fucosylation at the N-linked N-acetyl glucosamine (GlcNAc). This N-glycan type cannot be released by PNGase F, serving as a negative control for the analysis^27^. Incubation of HRP with PNGase Rc for 30 min resulted in almost complete deglycosylation, showing a clear band shift toward the deglycosylated protein (∼33.9 kDa) (**Figure 3a**, **I**) and thereby confirming the ability of PNGase Rc to release α-1,3-core fucosylated plant N-glycans, as previously shown for other members of the acidic PNGases^37, 42^.

The second investigated protein, RNase B (∼17 kDa), is glycosylated by a single, high mannose type N-glycan (Man_5-9_GlcNAc_2_)^43^. Here, deglycosylation with PNGase Rc was clearly visible after overnight incubation and revealed a new protein band consistent with the molecular weight of the unglycosylated protein (∼14 kDa). The corresponding band after using PNGase F showed a much lower band intensity (**Figure 3a, II**). From these results, we concluded that PNGase Rc released N-glycans with high mannose content more efficiently than PNGase F. We then examined the substrate fetuin (48.4 kDa) from fetal calf serum, with three N-, and four O-linked glycans having a high degree of sialylation^44^. Notably, the release of glycans by any PNGase is limited to N-linked glycans only. Again, PNGase Rc showed the onset of deglycosylation within 30 min, resulting in the disappearance of the initial fetuin band corresponding to N- and O-glycosylated fetuin (**Figure 3a III**), whereas a comparable release of N-glycans with PNGase F was observed only after six hours.

As a final example, we compared the efficiency of PNGase Rc and PNGase F in deglycosylation of the multiply glycosylated extracellular domain (ECD) of human signal-regulating protein alpha (SIRPα) using MS. Our results showed that upon overnight incubation with PNGase Rc at 37 °C and pH 2.5 all N-glycans were released (**Figure 3b, I**), whereas an identical incubation time with PNGase F at pH 7.5 resulted in an incompletely deglycosylated protein with non-resolvable mass spectra data (**Figure 3b, II**)

### Implementation of PNGase Rc for deglycosylation in a bottom-up HDX-MS workflow

To analyze the performance of the novel PNGase Rc for peptide deglycosylation under harsh HDX quench conditions requiring low temperature, low pH and short incubation times, we used the glycopeptide *EEQY**N**STYR* derived from a tryptic digest of trastuzumab. Thus, we incubated the glycopeptide at pH 2.5 in a water-ice bath (∼0 °C) for 2 minutes and relatively quantified the areas of the extracted ion chromatogram peaks (XICs, **Figure S2; Table S1**) of the glycosylated and deglycosylated peptide (**Figure 4a**). To determine an optimal E:S ratio resulting in complete peptide deglycosylation, we performed the assay using increasing concentrations of PNGase Rc. Our results showed that an E:S ratio of approximately 1:2.6 resulted in full deglycosylation of the peptide (**Figure 4b**). We next used this E:S ratio and added increasing concentrations of TCEP, urea or GdmCl to evaluate the deglycosylation activity of the PNGase Rc under harsh denaturing and reducing conditions. Our results revealed a most efficient N-glycan release in the presence of up to 200 mM TCEP and up to 4 M urea (**Figure 4c**). However, the addition of 1 M GdmCl decreased the activity by ∼ 50%, whereas higher concentrations of GdmCl (2 - 4 M) completely abolished the enzyme function (**Figure 4c**).

**Figure 4.**
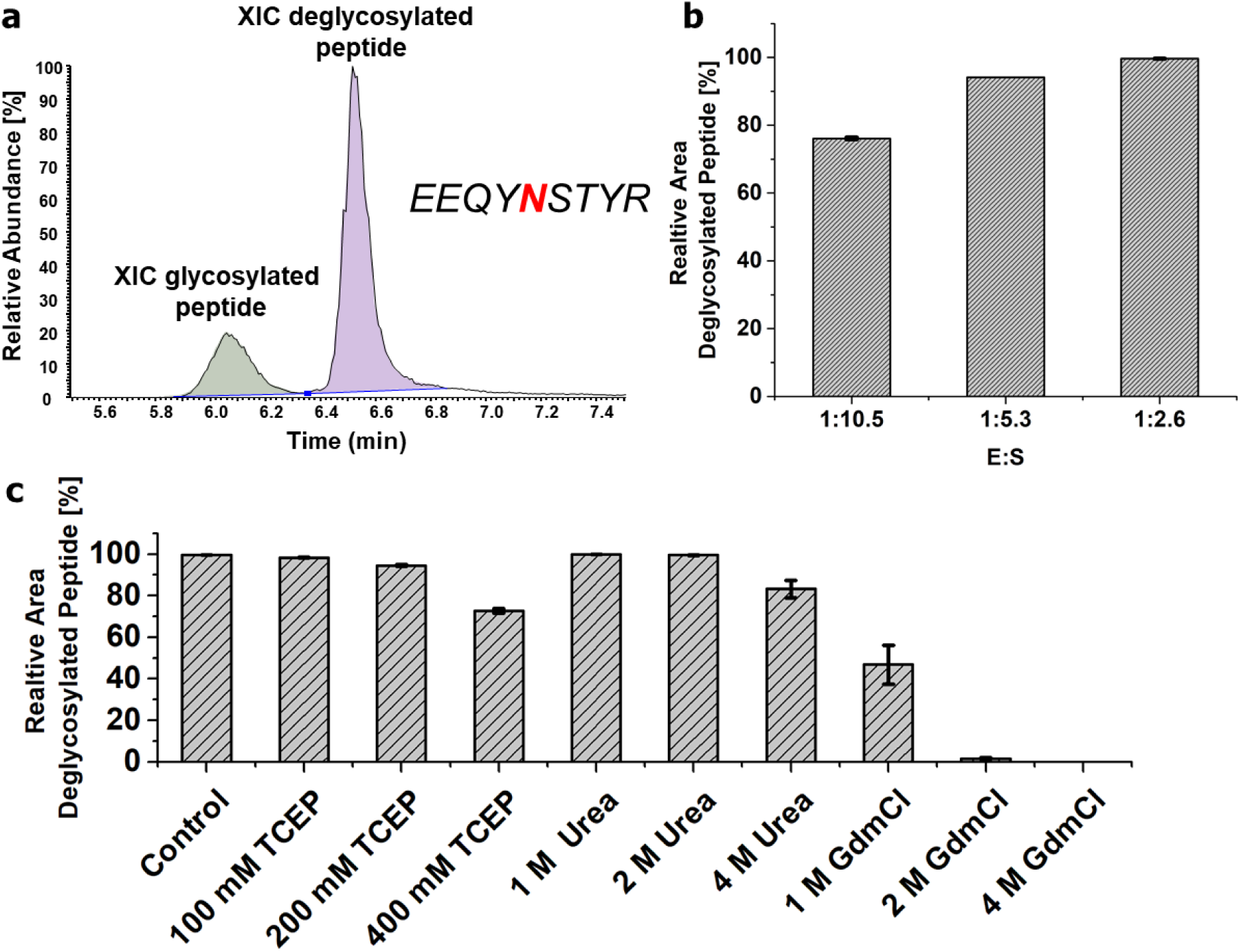
Influence of various of TCEP, urea and GdmCl concentrations on peptide deglycosylation by PNGase Rc under HDX quenching conditions. (**a**) XIC of the deglycosylated and glycosylated EEQYNSTYR peptide species for peptide-based deglycosylation assay (pH = 2.5, t = 2 min, T = 0 °C). (**b**) Deglycosylation activity using different E:S ratios: the deglycosylated peptide area was monitored relative to the sum of glycosylated and deglycosylated peptide peak areas). (**c**) Enzymatic activity of PNGase Rc in the presence of increasing concentrations of TCEP, urea and GdmCl using an E:S of 1:2.6.

### Implementation of PNGase Rc in a bottom-up HDX-MS workflow for epitope mapping

We continued and assessed the applicability of the novel PNGase Rc in a bottom-up HDX-MS workflow to determine the epitope of a single domain antibody (nanobody, Nb) targeting the extracellular domain (ECD) of human SIRPα (Wagner *et al.*, manuscript in preparation). The ECD of human SIRPα can be considered a challenging target as it comprises three disulfide bonds and five predicted N-glycosylation sites (**Figure 5a**). In a first step, we determined the N-glycosylation profile using tryptic digestion followed by tandem mass spectrometry (LC-MS/MS) to evaluate the micro-, and macroheterogeneity (**Figure S3**). The analysis revealed a high microheterogeneity for most of the N-glycosylation sites. Whereas the N240 and N262 are almost fully occupied with N-glycans, we found more than 10% of N215, more than 30% of N80 and 90% of N289 non-occupied with N-glycans. Next, we performed HDX-MS peptide analysis of SIRPα using an in-solution proteolysis step after deuterium labeling with bead-immobilized pepsin. To achieve high peptide coverage, peptic digestion was performed under reducing and denaturing conditions in the presence of 100 mM TCEP and 2 M GdmCl, resulting in the identification of 107 peptides (81% sequence coverage). Identification of cysteine-containing peptides showed successful reduction and hydrolysis of disulfide bonds under the applied conditions. However, our analysis revealed a low sequence coverage in proximity to the N-glycosylation sites. To improve this, we performed post-proteolytic deglycosylation using the PNGase Rc. Upon incubation with the enzyme under HDX quench conditions and in the presence denaturing and reducing agents, we identified another 7 peptides spanning N-glycan motifs and achieved an improved sequence coverage of 86% (**Figure 5**). Especially the sequence coverage and peptide redundancy in the regions encompassing N-glycosylated asparagine N215 and N240 benefited from the additional deglycosylation step. Both residues are almost fully glycosylated (**Figure S3**). Additionally, we identified two peptides including N215 and a previously inaccessible pepsin cleavage site. Furthermore, deglycosylated peptides covering N80 were found in three times higher signal intensity compared to their non-glycosylated peptide analogs (**Figure S4**), which are chromatographically separated from each other. Consequently, more reliable HDX information was obtained for the region near N80 as well.

**Figure 5.**
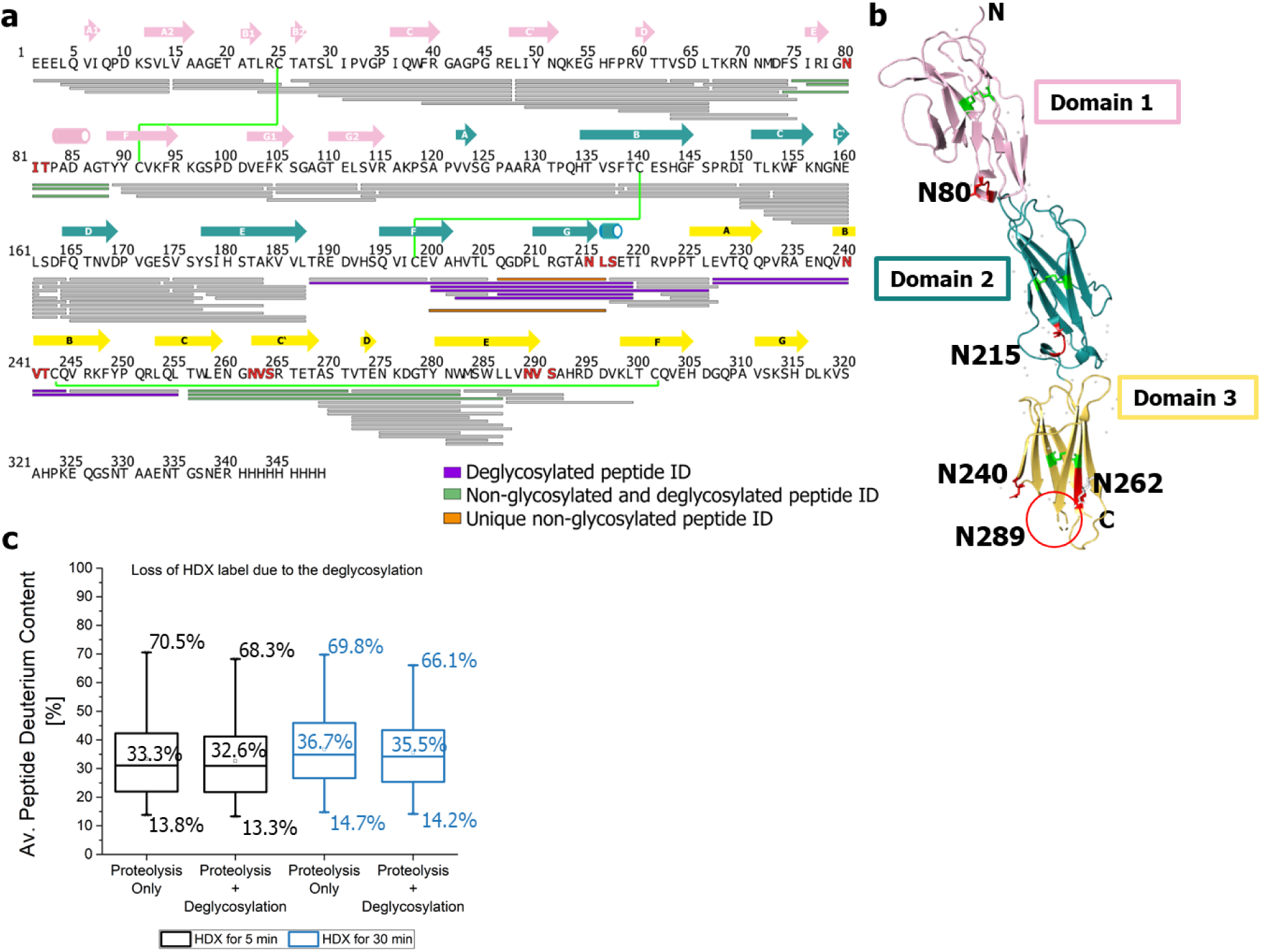
Peptide coverage map of human SIRPα with and without additional deglycosylation, x-ray crystal structure (PDBe: code: 2wng^45^), and loss of the HDX label due to additional deglycosylation. (**a**) Sequence coverage and redundancy of SIRPα proteolyzed for 2 min at ∼0 °C with and without additional deglycosylation. Peptides identified with and without additional deglycosylation: grey; new peptide identifications upon deglycosylation: purple; peptides identified with increasing signal intensity due to the deglycosylation of the partially occupied N-glycosylation site: light green; new peptides with previously inaccessible pepsin cleavage site: orange; disulfide bonds: green lines linking the involved cysteins, N-glycosylated asparagine residues: red. Amino acids involved in a secondary structure are shown with arrows (β-sheet) and tubes (α-helix) according to the structure^45^. (**b**) X-ray structure of SIRPα with predicted N-glycosylation (red) and disulfide bonds (light green). (**c**) Average HDX data of 90 peptic peptides obtained from SIRPα proteolysis determined in independent triplicates with and without additional deglycosylation. The average and minimal/maximal HDX values are shown.

Using a mixture of fully deuterium labeled model peptides, an average HD back exchange of ∼25% was determined for the HDX-MS system and workflow (**Figure S5).** We also assessed the potential increase in back exchange due to the deglycosylation of SIRPα by comparing 90 peptic peptides labeled for 5 and 30 min in D_2_O in triplicate analysis with and without the additional post-HDX deglycosylation protocol (**Figure 5c**). The results show that additional deglycosylation causes a minor decrease of the average peptide HDX of ∼0.7 and ∼1.2% after 5 and 30 min labeling, respectively. Thus, we conclude that the slight increase in back-exchange is an acceptable trade-off compared to the ability to obtain additional sequence coverage and redundancy and thus extract reliable HDX data and detect potential epitopes near N-glycan moieties.

Next, the deglycosylation workflow was evaluated for epitope mapping of a Nb, which was recently identified as a high affinity binder of SIRPα (Wagner *et al*., manuscript in preparation). Therefore, differential HDX experiments were performed for 0.5, 5, and 30 min in triplicate and for 500 min in duplicate. In total, we extracted HDX information for 101 peptides, corresponding to a sequence coverage of 86% (**Figure 6a**). Based on the variance of the replicate measurements, the significance threshold for the HDX differences was 0.34 Da at a 95% confidence level. Significant deuterium reduction upon Nb binding was observed in domain 2 of SIRPα (**Figure 6**). Several peptides spanning amino acid residues (aa) 190-225 and thus the N-glycosylated residue N215 showed significant HDX reduction upon binding (**Figure 6, Figure S6**). This suggests that the binding to the Nb causes a stabilization of the structure in this region. By extracting HDX data from overlapping peptide residues, we were able to narrow down the protected region. No significant difference in HDX of peptides in the N-terminal to region (aa 1-200) and C-terminal region (aa 220-349) was observed (**Figure 6b**). Deuteration for ≥30 min led to a disappearance of the HDX difference, which might originate from a dynamic interaction of the Nb with an amino acid loop subjected to high intrinsic deuteration. Notably, with the additional deglycosylation step increased the number of identifiable peptides for HDX analysis in this region by 42%, resulting in higher redundancy and thus higher confidence in the extracted HDX data.

**Figure 6.**
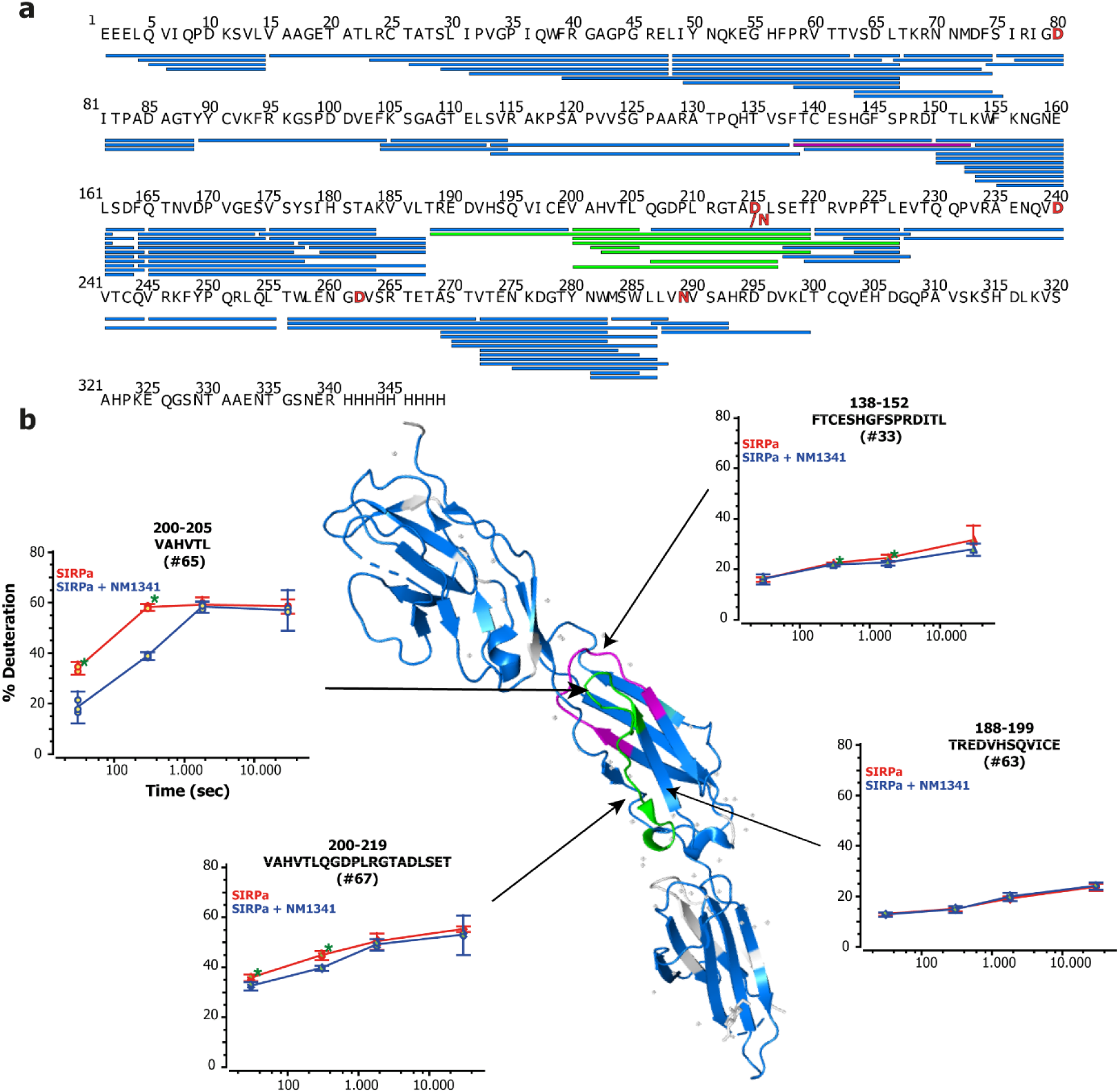
Differences in HDX due to the Nb mapped on the structure model of SIRPα (PDB: 2wng). (**a**) Sequence coverage map of SIRPα with peptides that show no significant changes (blue) and peptides with highly (green) and minor but significant deuteration differences (pink). (**b**) Crystal structure of SIRPα with protected region colored according to (**a**). Grey areas denote the absence of HDX information. HDX uptake plots of peptides corresponding to aa 138-152, 188-199, 200-205 and 200-219 of SIRPα in presence (blue) and absence (red) of the Nb. Error bars are based on the 95 % confidence level. Significant difference on basis of the Student’s t-test (p≤0.05) are labeled with an asterisk. Uptake plots of all analyzed peptides are listed in **Figure S5**

A minor but significant HDX difference was observed for the peptide corresponding to aa 138-152, while the peptide corresponding to aa 138-149 showed no significant difference upon Nb binding. However, this region is in proximity to aa 190-225 in the three-dimensional structure and might contribute to the binding interface of the Nb:SIRPα complex (**Figure 6b**). According to the consensus guidelines^11^, a summary of HDX-MS parameters (**Table S2**), HDX uptake plots of the differential analysis for all peptides with and without the Nb (**Figure S6**) as well as a summary table of the HDX uptake data (**Table S4**) can be found in the supplementary data. Based on the observation that the determined binding interface of the Nb is in close proximity to the N-glycan attached to N215, which might result in an altered binding behavior depending on the glycosylation state, we finally determined the binding affinity of Nb for the glycosylated and deglycosylated protein by biolayer interferometry (BLI) (**Figure S7**). For this, we deglycosylated SIRPα in its native protein conformation and compared the affinity constants of the Nb to the deglycosylated and glycosylated protein. No significant differences were found indicating that N-glycosylation does not influence the Nb-SIRPα interaction (**Figure S7**). In summary, our results show how the use of novel PNGase Rc improved the HDX-MS data for the elucidation of epitopes in a highly glycosylated and disulfide bonded protein.

## Discussion

Peptide N-glycanases are indispensable for characterizing the significance of sugar structures of proteins by means of enzymatic hydrolysis of N-glycans or for making proteins accessible to mass spectrometric and structural analysis^10, 28, 46^. Regarding their substrate specificity, peptide N-glycanases differ both in terms of the N-glycans they can hydrolyze and in terms of protein conformation susceptibility^47^. As for glycan specificity, hydrolysis is particularly influenced by the nature of fucose linkage to the core region. Of the two commercially available and widely used PNGases - PNGase F and PNGase A^46, 48^ - only PNGase A can hydrolyze N-glycans with both, the α(1,3)- and the α(1,6) core fucosylation, while PNGase F only releases N-glycan moieties with α(1,6) core fucosylation^27^. Likewise, both PNGases differ in their optimal ambient conditions. While PNGase F has an optimal pH range of 6.0 - 9.5^29^, PNGase A preferably works under slightly acidic (pH 4.0 - 6.0) conditions^30^. Whereas PNGase F is readily expressed in standard bacterial systems such as *E. coli*, the expression and purification of PNGase A, which itself comprises twelve potential N-glycosylation sites^27^, is not possible in simple prokaryotes^27, 37^. More recently, two variants, each isolated from *Terriglobus roseus* (PNGase Tr) and *Dyella japonica* (PNGase Dj), that both showed improved properties regarding their specificity and working conditions have been described^37, 42^. However, their availability is still very limited, especially since they were not yet produced on a larger scale in simple expression systems.

With the PNGase Rc derived from the soil bacterium *Rudaea cellulosilytica*, here we present a novel PNGase which shows <50% sequence homology as determined by the BLASTp algorithm to the other recently published acidic PNGases (**Table S3**). Following heterologous expression, PNGase Rc can be easily obtained from *E.coli* in large quantities and sufficient purity. With respect to its substrate specificity, our data showed that the novel PNGase Rc is largely insensitive to the protein conformations. This enables deglycosylation of native, multiply glycosylated proteins and opens up new applications, such as deglycosylation of proteins prior to X-ray crystallography or investigation of the influence of N-glycans on protein-protein binding in biological assays. Likewise, the enzyme is able to hydrolyze N-glycans with both possible linkages of fucose to the glycan core region. Thus, it addresses sugar structures as they occur in plants, invertebrates and mammals^49, 50^.

Considering that the different variants of PNGases differ in size, number of disulfide bridges, and their own glycosylation, they have different sensitivities to reducing and denaturing substances. With the PNGase Rc we selected a very stable variant which tolerates the presence of reducing agents under HDX quench conditions, such as 0.4 M TCEP or 1 M GdmCl and retains ∼80% of its activity even in the presence of 4 M urea. Thus, it showed a higher stability and activity under harsh buffer conditions compared to the commercially available PNGase A^33^. Notably, its sensitivity to detergents remains to be verified.

Due to its improved pH optimum, stability, activity and broad specificity, PNGase Rc is not only suitable for the analysis of intact native proteins, but especially for the analysis of protein interactions by HDX-MS, which we demonstrated by elucidating the binding region of a Nb to a multiple N-glycosylated antigen. Here, the use of the enzyme led to a significant increase in sequence coverage and redundancy of the peptides obtained, particularly in the region of the fully occupied N-glycosylation sites.

In summary, PNGase Rc will open new opportunities for the study of glycosylated proteins under a variety of conditions due to its advantageous properties. For future applications, the enzyme could be immobilized on chromatographic support material, which would allow a highly efficient use in HDX applications with online pepsin digestion prior to chromatographic separation.

## Experimental Section

### Proteins

Trastuzumab was kindly donated by MAB Discovery GmbH (Polling, Germany), fetuin from fetal calf serum (New England Biolabs, Ipswich MA, USA), RNase B from bovine pancreas (New England Biolabs, Ipswich MA, USA), and horse radish peroxidase (HRP, Sigma-Aldrich, Munich, Germany), PNGase F from *F. meningosepticum* (R&D Systems, Wiesbaden, Germany), human SIRPα (Glu 31 - Arg 370 (ACROBiosystems, Newark, DE, USA)

### PNGase Rc expression and purification

See the Supporting Information

### Activity determination using antibody deglycosylation and LC-MS analysis

See the Supporting Information

### Intact deglycosylation of HRP, RNase B, fetuin and SIRPα monitored by SDS-PAGE and MS

Native protein deglycosylation of 20 µL (2 mg/mL) stock solutions of HRP (45.5 µM), fetuin (41.3 µM) or RNase B (117.7 µM) were carried out by adding 70 µL 100 mM citrate-NaOH buffer pH 3.5 or 100 mM Tris HCl pH 7.5 for PNGase Rc or F, respectively. Deglycosylation was initiated by adding 10 µL PNGase F or Rc for HRP and RNase B or 9 µL for fetuin, respectively achieving an enzyme substrate (E:S) ratio of ∼1:48 (M:M). After incubation for 0.5, 1, 3, 6 and 23 h at 37 °C, aliquots of 19.5 µL were quenched by adding 7.5 µL 4 X LDS sample buffer (Life Technologies, Carlsbad, CA, USA), 3 µL 10X sample reducing agent (Life Technologies, Carlsbad, CA, USA) and heat-denaturation for 5 min at 95 °C. The acidic pH of the samples prepared with PNGase Rc was neutralized by adding 10% ammonia prior to analysis. Gel electrophoresis was performed using the commercial NuPAGE® system (Life Technologies, Carlsbad, CA, USA) with a 4–12%, 12 Well, Bis-Tris gel according to the manufacturer’s protocol. SeeBlue^TM^ Plus2 pre-stained standard (Life Technologies, Carlsbad, CA, USA) was used as molecular marker (5 µL) and 15 µL of each sample was loaded onto the gel. A negative control was prepared using the buffer system and protocol of PNGase Rc incubated for 23 h with water instead of the protein. Protein bands were visualized using Blue BANDit^TM^ protein stain (VWR Life Science, Darmstadt, Germany).

For intact SIRPα deglycosylation 2 µL (40.2 µM) sample was diluted 1:10 either with PBS pH 7.4 (PNGase F) or 100 mM citrate-NaOH buffer pH 2.5 (PNGase Rc). Deglycosylation was initiated by adding 0.5 µL, 3.6 µM PNGase to achieve a E:S ratio of 1:45 (M:M). Samples were incubated overnight at 37 °C and analyzed subsequently using the intact protein HPLC-MS analysis described above injecting 1.2 µg SIRPα.

### Analysis of SIRPα N-glycosylation microheterogeneity

See the Supporting Information

### Peptide deglycosylation assay to test TCEP, GdmCl and Urea tolerance

A tryptic proteolysis of trastuzumab was performed and purified as previously described by Zeck *et al*.^51^. Peptide deglycosylation was performed in a 40 µL sample volume consisting of 10 µL (∼2.62 µM) of the tryptic trastuzumab digest, 5 µl of 1, 2 or 4 µM PNGase Rc in 100 mM citrate-NaOH buffer pH 2.5, 4 µL 1 M citrate-NaOH buffer pH 2.5, 1 µL H_2_O and 20 µL 100 mM citrate-NaOH buffer pH 2.5. Stock solutions of 8 M GdmCl and urea were prepared in 100 mM ammonium formate solution pH 2.5. A stock solution of 1 M TCEP in H_2_O was prepared and basified with ammonia to pH 2.5. The PNGase Rc tolerance against denaturing and reducing agents was probed using the above described assay. Thus, 5 µL 4 µM PNGase Rc was used and 20 µL100 mM citrate-NaOH buffer pH 2.5 was replaced with varying concentration of the stock solutions of TCEP, urea or GdmCl with final concentrations of 100 - 400 mM, 1 - 4 M and 1 – 4 M, respectively.

Samples were incubated for 2 min in a water-ice bath (∼0 °C) and subsequently were snap frozen and stored at -80 °C until analysis by HPLC-MS. After thawing immediately 1 µL of each sample was injected onto the HPLC-MS system. Samples were analyzed by HPLC-MS using an Orbitrap Eclipse^TM^ Tribrid^TM^ instrument coupled to an UltiMate^TM^ 3000 RSLCnano system (Thermo Fisher Scientific, Dreieich, Germany). For detailed method description, see supporting information.

### HDX peptide list generation of SIRPα with and without post-proteolysis deglycosylation by PNGase Rc

SIRPα was buffer exchanged to PBS pH 7.4 using sing Zeba^TM^ Spin Desalting Columns 7K MWCO according to the manufacturer’s protocol. SIRPα was incubated 1:0.5 (v/v) in PBS (pH 7.4) and equilibrated for 10 min at 25 °C. Samples were diluted 1:10 (v/v) in PBS (pH 7.4) incubated and 15 µL aliquots were quenched 1:2 (v/v) with ice-cold quenching solution (200 mM TCEP with 1.5% formic acid (v/v) and 4 M GdmCl in 100 mM ammonium formate solution pH 2.2) resulting in a final pH of 2.5. Quenched aliquots were immediately snap frozen in liquid nitrogen and stored at – 80 °C. Settled gel of immobilized pepsin (Thermo Scientific, Schwerte, Germany) was prepared by centrifugation of 60 μL 50% slurry (in ammonium formate solution pH 2.5) for 3 min at 1,000 x g and 0 °C. The supernatant was discarded, protein aliquots were thawn, added to the settled pepsin gel and digested for 2 min in a water-ice bath. Immobilized pepsin was removed by centrifugation at 1,000 x g for 30 sec at 0 °C using a 0.65 µm filter inlet (Merck Millipore, Darmstadt, Germany) and the flow through was injected immediately into the HPLC-MS/MS system. Peptides were detected by tandem MS as described previously^52^. A VanGuard column (2.1 x 5 mm, particle: 1.7 µm; 130 Å; Waters GmbH, Eschborn, Germany) was used in order to protect the analytical column from PNGase Rc and undigested protein fragments. Notably, peptide identification was improved by alternative proteolysis for 5 min at 0 °C and using a quenching buffer containing 400 mM and 4 M GdmCl. Additional deglycosylation was conducted using 5 µL (4 µM) PNGase Rc in 100 mM ammonium formate solution pH 2.5 placed underneath the 0.65 µm filter inlet (**Figure S8**) according to Jensen *et al*. ^33^. Deglycosylation was initiated by centrifugation (1,000 x g, 30 sec, 0 °C) and samples were incubated for 2 min at 0 °C. HPLC-MSMS peptide analysis was performed as previously described^52^. For detection of deglycosylated peptide species, deamidation at asparagine residues was added as dynamic modification.

### HDX-MS Nb epitope mapping of SIRPα with post-proteolysis deglycosylation

For HDX analysis SIRPα (42 µM) was either incubated 1:0.5 (v/v) with PBS (pH 7.4) or Nb (97.36 µM, in PBS) and was incubated for 10 min at 25 °C. HDX of the pre-incubated complex was initiated by diluting 1:10 (v/v) in deuterated PBS (pH 7.4). Using the equations of Kochert *et al*. ^53^ and the K_D_ of 32.4 nM, 95% of SIRPα is theoretically bound to the Nb during HDX labeling. After incubation for 0.5, 5 and 30 and 500 min aliquots of 15 µL were quenched by adding 1:2 (v/v) with ice-cold quenching solution (200 mM TCEP with 1.5% formic acid (v/v) and 4 M GdmCl in 100 mM ammonium formate solution pH 2.2) resulting in a pH of 2.5. Samples were immediately snap frozen using liquid nitrogen and stored at -80 °C. The above-described 2 min peptic proteolysis with subsequently 2 min deglycosylation was applied. Peptides were separated by RP-HPLC and analyzed in MS1 mode as previously described by Wagner *et al*. ^52^. For examination of the additional deuterium loss due to deglycosylation, the protocol described above was performed with and without the additional deglycosylation step. Thus, SIRPα was deuterated for 5 and 30 min and injected immediately after the 2 min proteolysis. Non-deuterated control samples were prepared with PBS prepared with H_2_O. Samples were prepared in independent triplicates and analyzed with HDExaminer v2.5.0 (Sierra Analytics, Modesto, CA, USA) using the centroid mass shift. Mapping of H/D exchange data on x-ray structures was conducted using PyMOL (v2.0.7 http://www.pymol.org)

### Binding kinetic determination by biolayer interferometry

See the Supporting Information

### Data availability

The data that support the findings of this study are available from the corresponding authors upon reasonable request.

### Authorship Contributions

M.G., J.V., S.M., D.S. U.R. and A.Z. designed the study; U.R., T.W., B.T. performed Nb selection and biochemical characterization; P.D.K expressed and purified the PNGase Rc; M.G. performed the PNGase activity, specificity and HDX-MS experiments; J.V. provided the PNGase sequence. M.G., P.D.K., A.Z., D.S., S.M. and U.R. analyzed data and performed statistical analysis. M.G. and A.Z. drafted the manuscript. All authors critically read the manuscript.

## Acknowledgments

This work received financial support from the State Ministry of Baden-Wuerttemberg for Economic Affairs, Labour and Tourism. The RSLC U3000 HPLC system and the Orbitrap Eclipse Tribrid Mass Spectrometer were funded from the European Regional Development Fund (ERDF).

## Supporting Information

### Supporting Experimental Section

#### PNGase Rc expression and purification

The nucleic acid coding for amino acids 17-581 of the putative PNGase homolog Rc (from Rudaea Cellulosilytica UniParc ID UPI00146EBC6F) was synthesized and ligated into a pET30a(+) *E.coli* expression vector by Genscript Ltd. (Nanjing, China). N-terminally His_6_-tagged PNGase Rc was expressed in *E.coli* strain BL21 (DE3) (New England Biolabs, Ipswich MA, USA). *E.coli* from 100 mL preculture grown in cultivation medium (Terrific Broth supplemented with 50 µg/mL Kanamycin) at 37 °C overnight were resuspended in 1 L fresh cultivation medium and grown until OD_600_ of 0,8 was reached. To induce protein expression 0,1 mM IPTG (isopropyl β-D-thiogalactopyranoside) was added and the culture was further incubated overnight at room temperature under constant stirring. Centrifuged *E.coli* pellet was resuspended in 30 mL 1 x PBS (phosphate buffered saline) supplemented with 363 mM NaCl, 50 mM Imidazole and 1 mM PMSF (phenylmethylsulfonyl fluoride). After one freeze thaw cycle and adding of Protease-Inhibitor Mix B, 60000 units lysozyme and 6000 Kunitz units DNAseI *E.coli* were disrupted by repeated pulsed sonication and end-over-end rotation for 90 min at 4°C. Lysate was centrifuged at 20000 x g and 4 °C. PNGase Rc was purified from the soluble extract by immobilized metal affinity chromatography using 1 mL HisTrap® FF (Cytiva, Marlborough, MA US) and ÄKTA pure 25 purification system (GE Healthcare, Uppsala, Sweden). Eluted protein peak was loaded on a HiLoad® 26/600 Superdex® 75 pg column (Cytiva, Marlborough, MA USA) for size exclusion chromatography using PBS as buffer. Purification of PNGAse Rc was examined by SDS-PAGE followed by Coomassie staining and western blotting. An anti-penta His antibody (QIAGEN, Hilden, Germany) and anti-mouse antibody labeled with Alexa 647 (Thermo Fisher Scientific, Waltham, USA) were used for detection. Protein purity was estimated via line scan of Coomassie stained SDS-PAGE using ImageJ software.

#### Activity determination using antibody deglycosylation and LC-MS analysis

For activity determination, 5 µL of 5 mg/mL (34.1 µM) trastuzumab in 20 mM histidine buffer (150 mM NaCl, pH 6.0) were diluted with 40 µL, 100 mM citrate-NaOH buffer pH 2.5 (for PNGase Rc) or 100 mM Tris HCl buffer pH 7.5 (for PNGase F). Deglycosylation was initiated by adding 5 µL of 0.78 µM PNGase Rc or PNGase F diluted in the corresponding buffer to achieve an enzyme:substrate (E:S) ratio of 1:44 (M:M). After incubation for 2.5, 5, 10, 20 and 40 min at 37 °C, aliquots of 10 µL were quenched using heat-denaturation at 95 °C for 5 min (PNGase Rc) or by adding 10 µL, 1% (v/v) formic acid (resulting in a pH = 2.3, PNGase F). Optimum temperature- and pH of PNGase Rc was determined using 3 µL of 5 mg/mL trastuzumab diluted in 22 µL buffer. Deglycosylation was initiated by adding 5 µL of PNGase Rc (0.47 µM). The temperature optimum was determined by incubation at 4 (on-ice), 20, 30, 37, 50, 60 and 70 °C in 100 mM citrate-NaOH buffer at pH 2.5. For determination of the pH-optimum, samples were prepared in 200 mM glycine HCl buffer at pH 2.0 or 100 mM citrate-NaOH buffer at pH 2.5, 3, 3.5, 4, 4.5 or 5 and were incubated at 37 °C. All samples were prepared in triplicates and quenched after 10 min by heat-denaturation for 5 min at 95 °C. Quenched samples were snap frozen in liquid nitrogen and stored at -80 °C until HPLC-MS analysis.Intact protein analysis performed on a QTOF-type mass spectrometer (MaXis HD UHR q-TOF, Bruker, Bremen, Germany) was conducted as described^16^. Data analysis was performed using Bruker Compass DataAnaylsis v5.3 software (Bruker Daltonik, Bremen, Germany). Charge deconvolution of the m/z spectra was performed with the MaxEnt deconvolution algorithm (Bruker Daltonic, Bremen, Germany).

#### Mass spectrometric analysis for peptide deglycosylation and inhibition by TCEP GdmCl, urea

Samples were analyzed by HPLC-MS using an Orbitrap EclipseTM TribridTM instrument coupled to an UltiMateTM 3000 RSLCnano system (Thermo Fisher Scientific, Dreieich, Germany) Injected samples were trapped on an AcclaimTM PepMapTM C18 (Thermo Fisher Scientific, Waltham, USA) (5 x 0.3 mm, 5 µm particle size, 100 Å pore size) at 120 µL/min flow rate for 0.25 min using 2% MeCN (v/v) + 0.05% trifluoroacetic acid (v/v, TFA) as solvent. Afterwards the injection valve was switched connecting the trapping column to the analytical column. Peptides were eluted from the trapping column and separated using an ACQUITY BEH M-Class C18 column, 150 x 0.075 mm, 1.7 µm particle size, 130 Å pore size (Waters GmbH, Eschborn, Germany). Nano HPLC-gradient parameters were used as follows: Eluent A (EA): 0.1% FA (v/v), eluent B (EB): 80% MeCN (v/v) + 0.1% FA (v/v). A linear gradient from 2-16% EB in 8 min was used at a flow rate of 0.6 µL/min followed by a wash saw tooth alternating from 2.5-99% EB. Total run time: 15 min. Column oven temperature: 50 °C. MS and MSMS analysis were performed with mass resolutions of120,000 and 30,000 resolution using the orbitrap mass analyser of the Orbitrap EclipseTM TribridTM instrument. Instrument configurations were used as follows: m/z: 200-2000; positive ion spray voltage of 2.1 kV and an ion transfer tube temperature of 275 °C.

### Analysis of SIRPα N-glycosylation microheterogeneity

15 µL SIRPα (1.5 mg/mL) was mixed with 56 µL denaturation buffer (400 mM Tris HCl, pH 8; 8 M GdmCl) and 5 µL 0.24 M dithiothreitol and incubated for 60 min at 37 °C. Afterwards 5 µL 0.6 M iodoacetamide was added and incubated for 15 min at room temperature. Buffer exchange was performed using Bio-Spin® P-6 gel Tris-buffer columns (RioRad Laboratories GmbH, Feldkirchen, Germany) as per the manufacturer’s instructions. Proteolysis was performed by adding 2 µL trypsin (0.25 µg/µL, E:S 1:50) overnight at 37 °C. Samples were analyzed on the same HPLC-MS system as described for the analysis of peptide deglycosylation and inhibition by TCEP, GdmCl, urea. Nano HPLC-gradient parameters were used as follows: Eluent A (EA): 0.1% FA (v/v), eluent B (EB): 80% MeCN (v/v) + 0.1% FA (v/v). A stepwise linear gradient from 2-16% EB in 8 min; 16-19% in 4 min; 19-29% in 8 min; 29-45% in 6.25 min. Flow rate: 0.6 µL/min. Additionally, washing with a saw tooth gradient alternating from 2-99% EB was applied. Total run time: 40 min. Column oven temperature: 50 °C. MS and MSMS analysis were performed with mass resolutions of120,000 and 30,000 resolution using the orbitrap mass analyser of the Orbitrap Eclipse^TM^ Tribrid^TM^ instrument. Samples were analyzed using BioPharma Finder software v4.1.53.14 (Thermo Fisher Scientific) with mass tolerances for MS = 6 ppm and MS/MS = 0.05 Da.

### Binding kinetic determination by biolayer interferometry

Binding affinity of the single domain antibody (nanobody, Nb) towards SIRPα (ACROBiosystems Cat.#SIA-H5225) was determined via biolayer interferometry (BLI) using Octet® RED96e system according to standard protocol. The Nb was biotinylated and immobilized on streptavidin biosensors (SA) (Sartorius Lab Instruments). SIRPα samples were incubated with PNGase Rc with an molar E:S ratio of 1:45 at 37 °C and pH 2.5 (100 mM Citrate NaOH) for 23 h. A control sample was prepared with buffer instead of the PNGase. Solution was adjusted to neutral pH by addition of Tris before BLI measurement. Dilution series ranging from 160 to 20 nM of SIRPα were measured and one reference was included per run. Additionally a non-treated control SIRPα was measured without pre-treatment in PBS. For affinity determination, the 1:1 global fit of the Octet® Data Analysis HT 12.0 software was used.

## Supporting Figures

**Figure S1:**
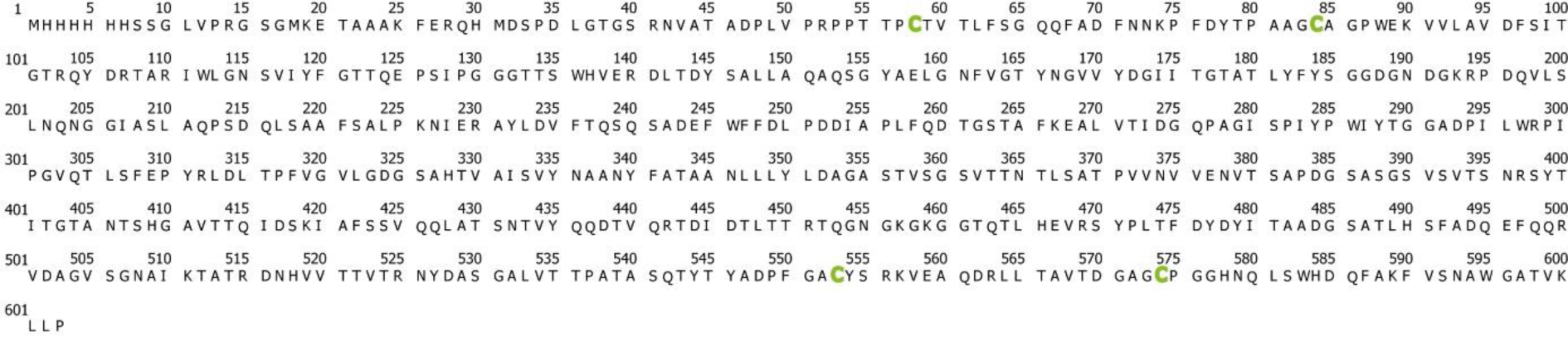
Sequence of the PNGase Rc construct. Theoretical mass = 64198.0 Da. cysteine residues are shown in green.

**Figure S2:**
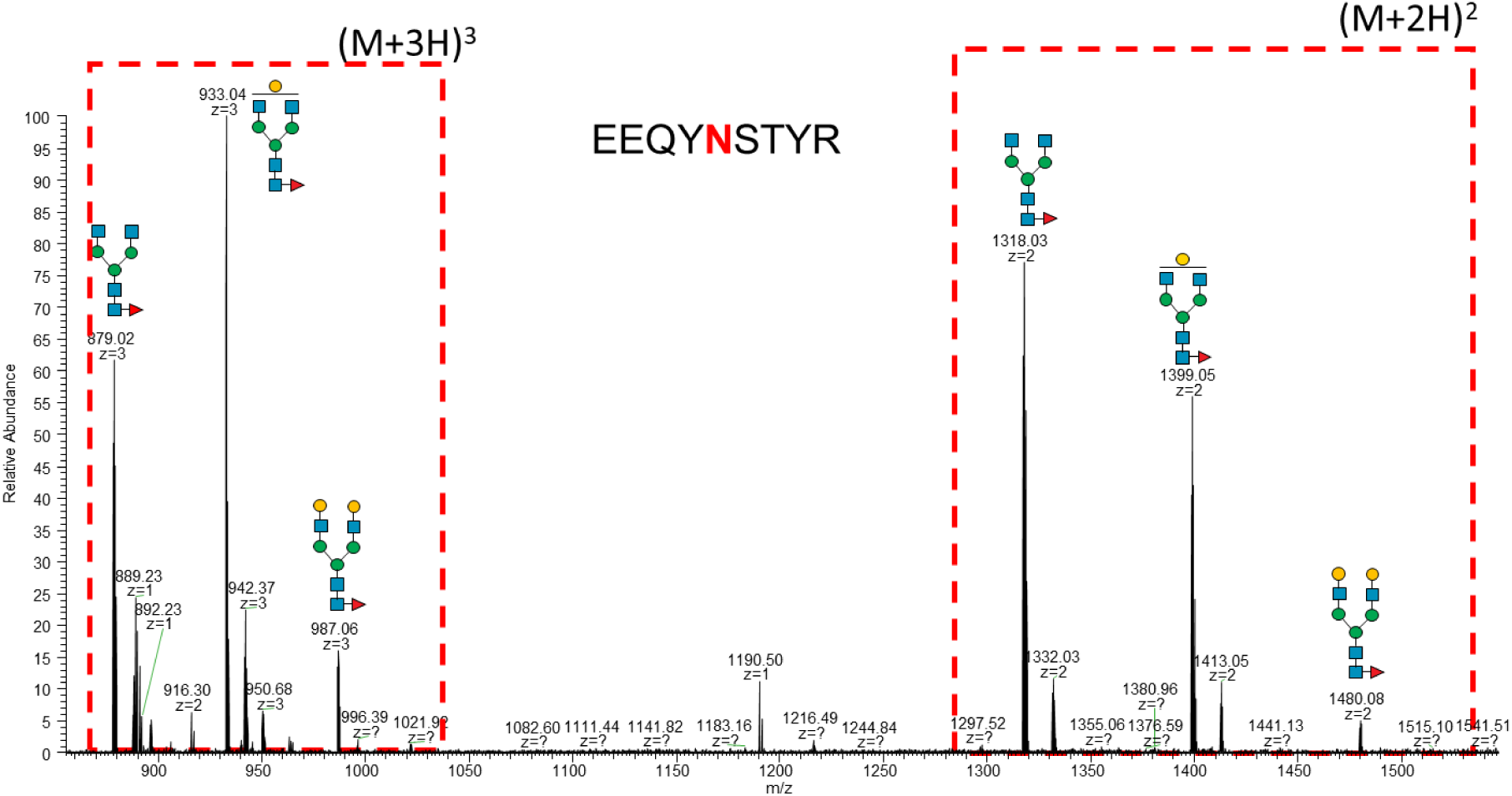
m/z spectrum summed up over retention time range from 5.8 to 6.5 min showing the *EEQYNSTYR* peptide from a tryptic trastuzumab digest with the different ion masses due to the peptide N-glycosylation.

**Figure S3:**
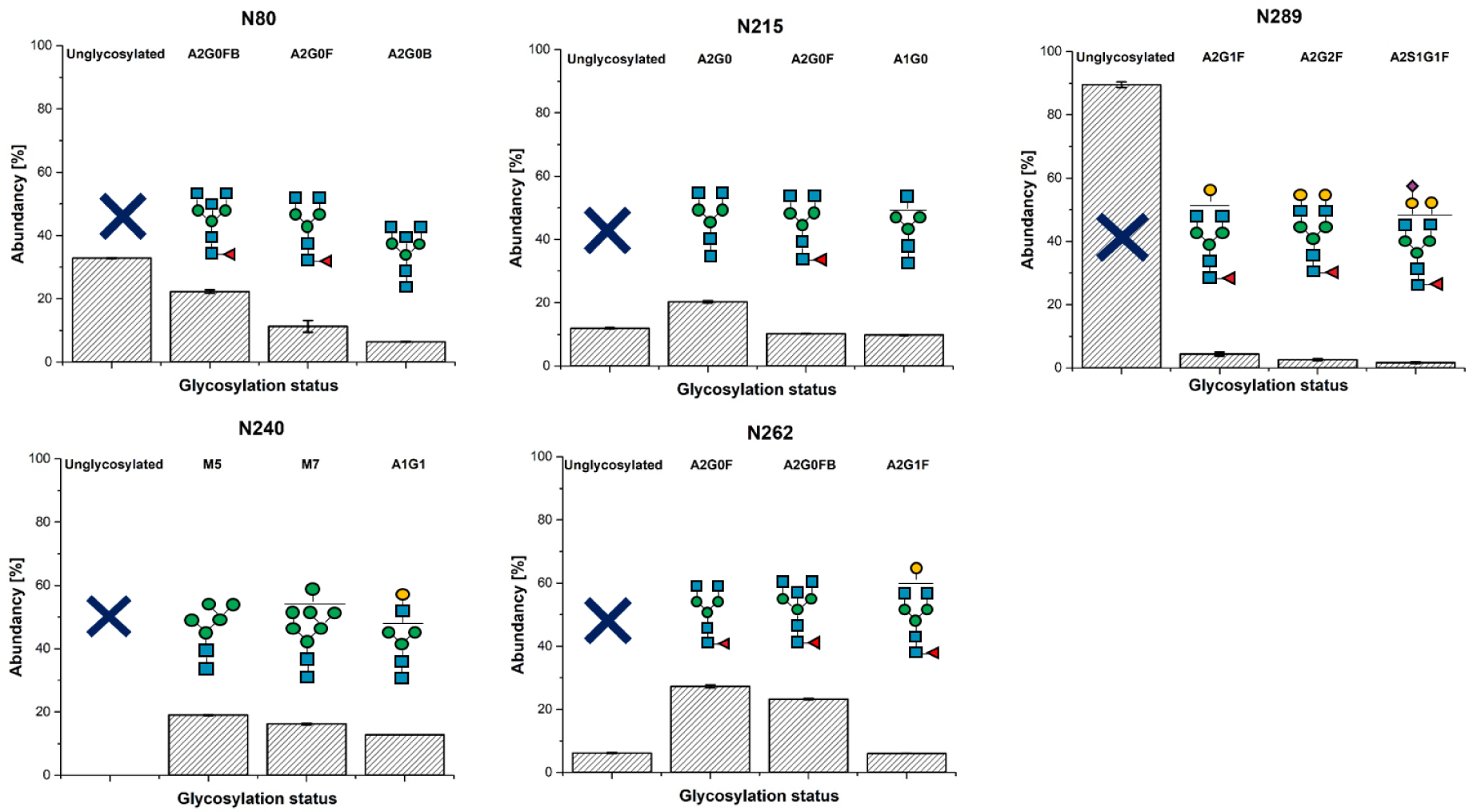
Determined N-glycosylation micro-, and macro heterogeneity of the five possible SIRPα N-glycan sites. SIRPα was proteolyzed overnight by trypsin. Peptides were identified by HCD fragmentation and by BioPharma Finder^TM^ v4.1.53.14 (Thermo Fisher Scientific). N-glycsoylation species are relatively quantified and the three most abundant N-glycans are shown.

**Figure S4:**
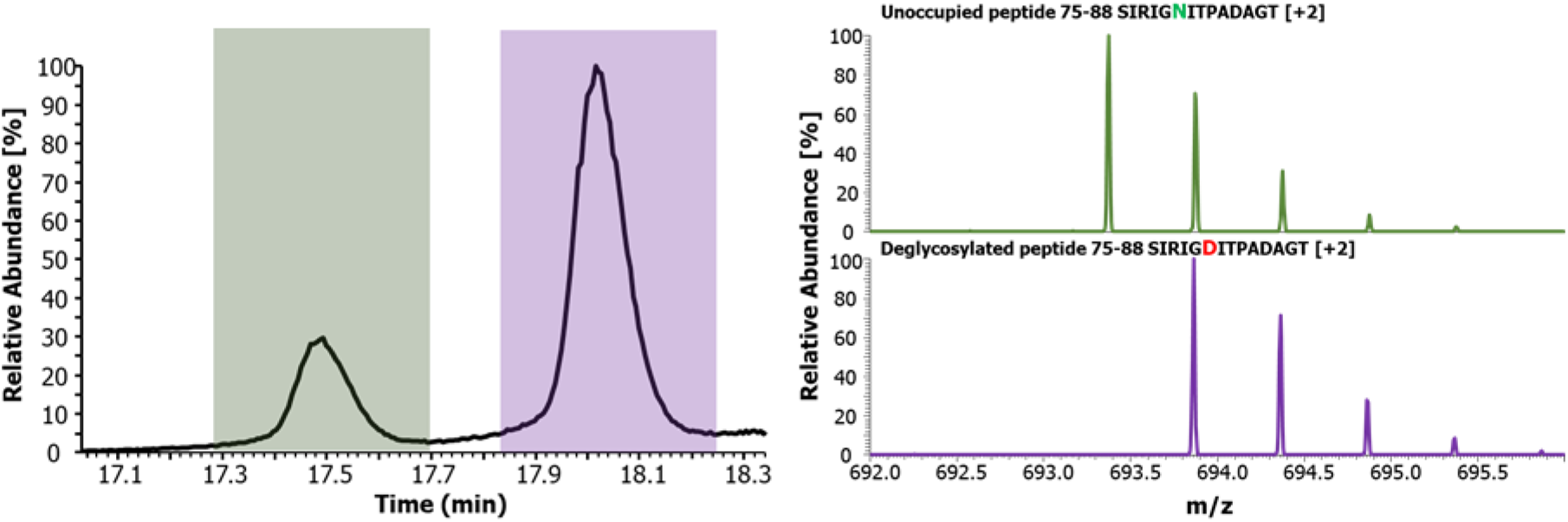
Extracted ion chromatogram (m/z 693.25-696.00) of the SIRPα peptide 75-88 (left) and m/z spectra summed up across the corresponding peaks (right). Deamidation to aspartic acid due to the deglycosylation by PNGase Rc of asparagine resulted in a mass shift of +0.98 Da.

**Figure S5:**
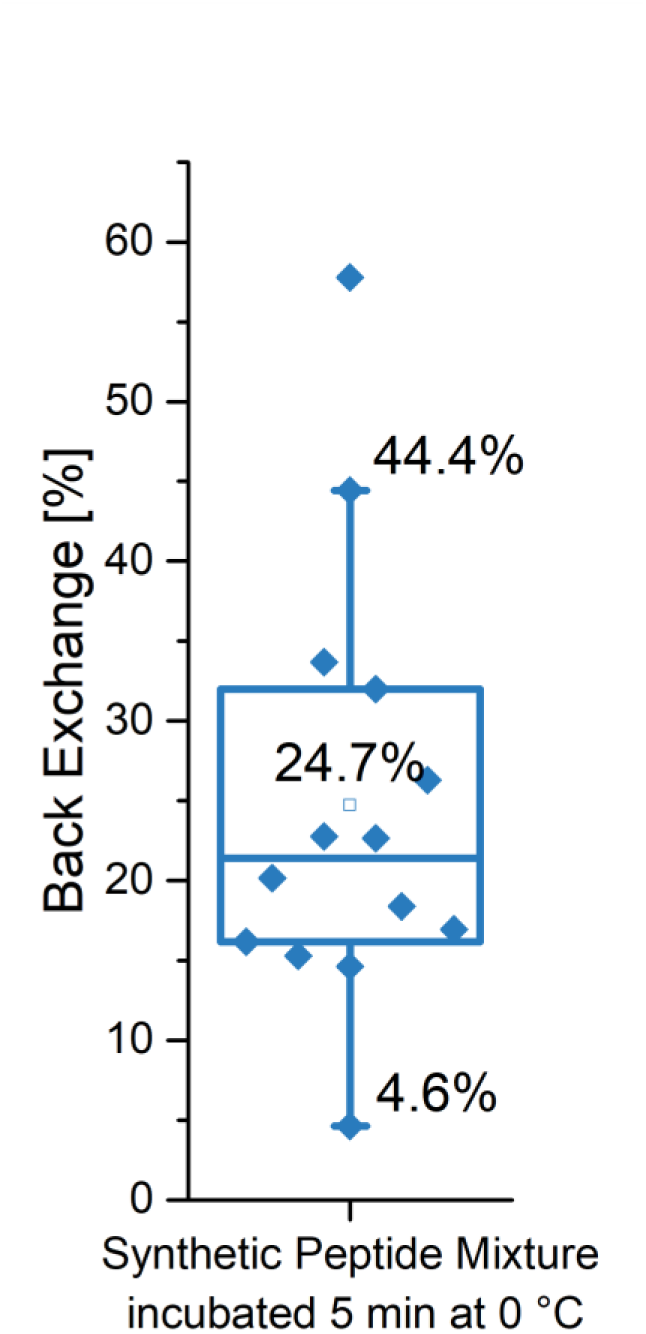
Determined back exchange using an artificial generated peptide mixture encompassing 14 synthetic peptides differing in their length and amino acid composition as described in^16^. Briefly, a peptides containing with 1 mg/mL of each peptide was diluted 1:10 in PBS and subsequently freeze-dried. Aliquots were restored 99.9% D2O and labeled overnight at 20 °C. Samples were quenched by adding 1:2 ice-cold quenching solution and a proteolysis step was simulated by storing the samples in a water-ice bath, 5 min prior to LC-MS analysis. HDX was analyzed in independent triplicates and the average back exchange is plotted.

**Figure S6.**
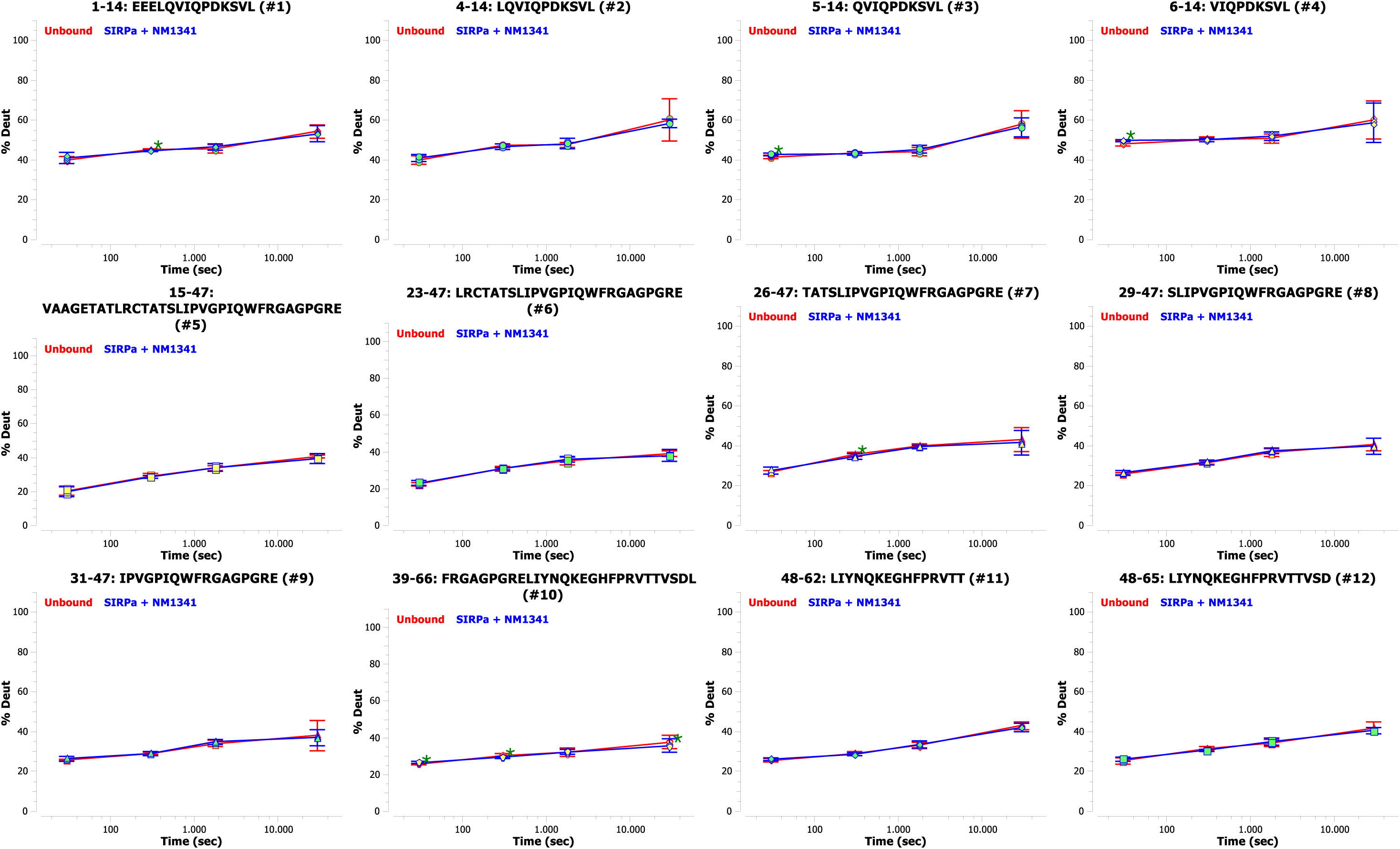
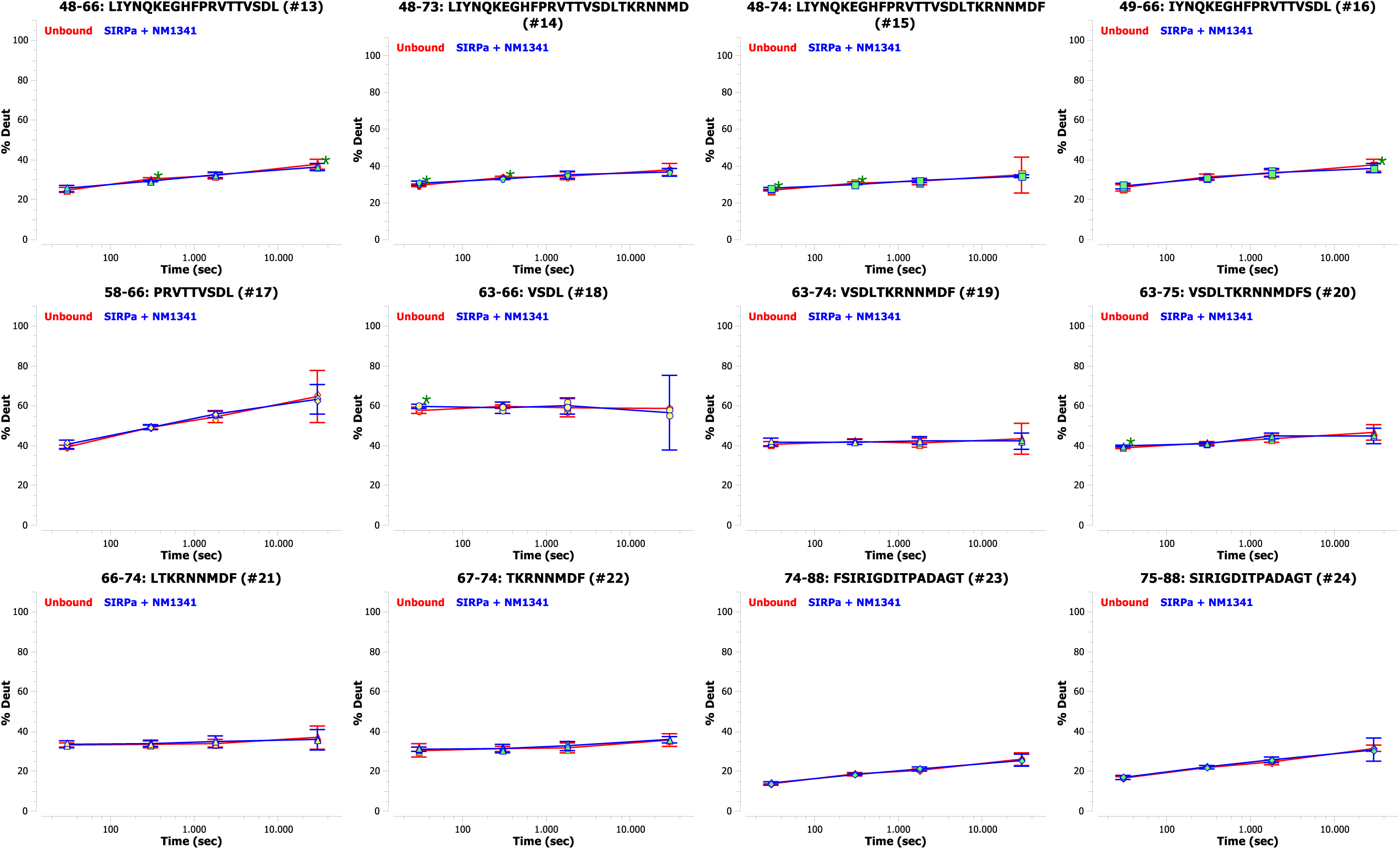
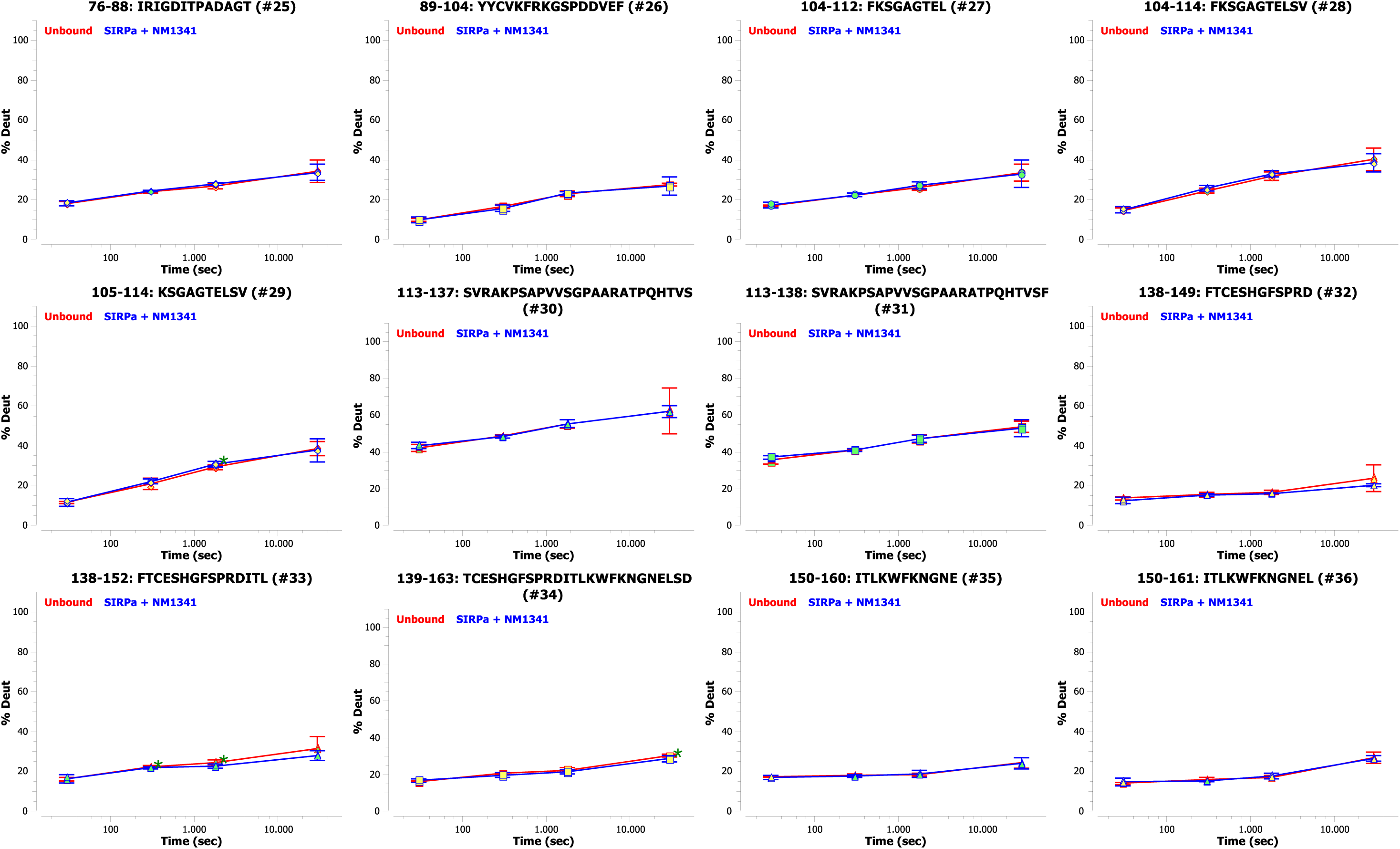
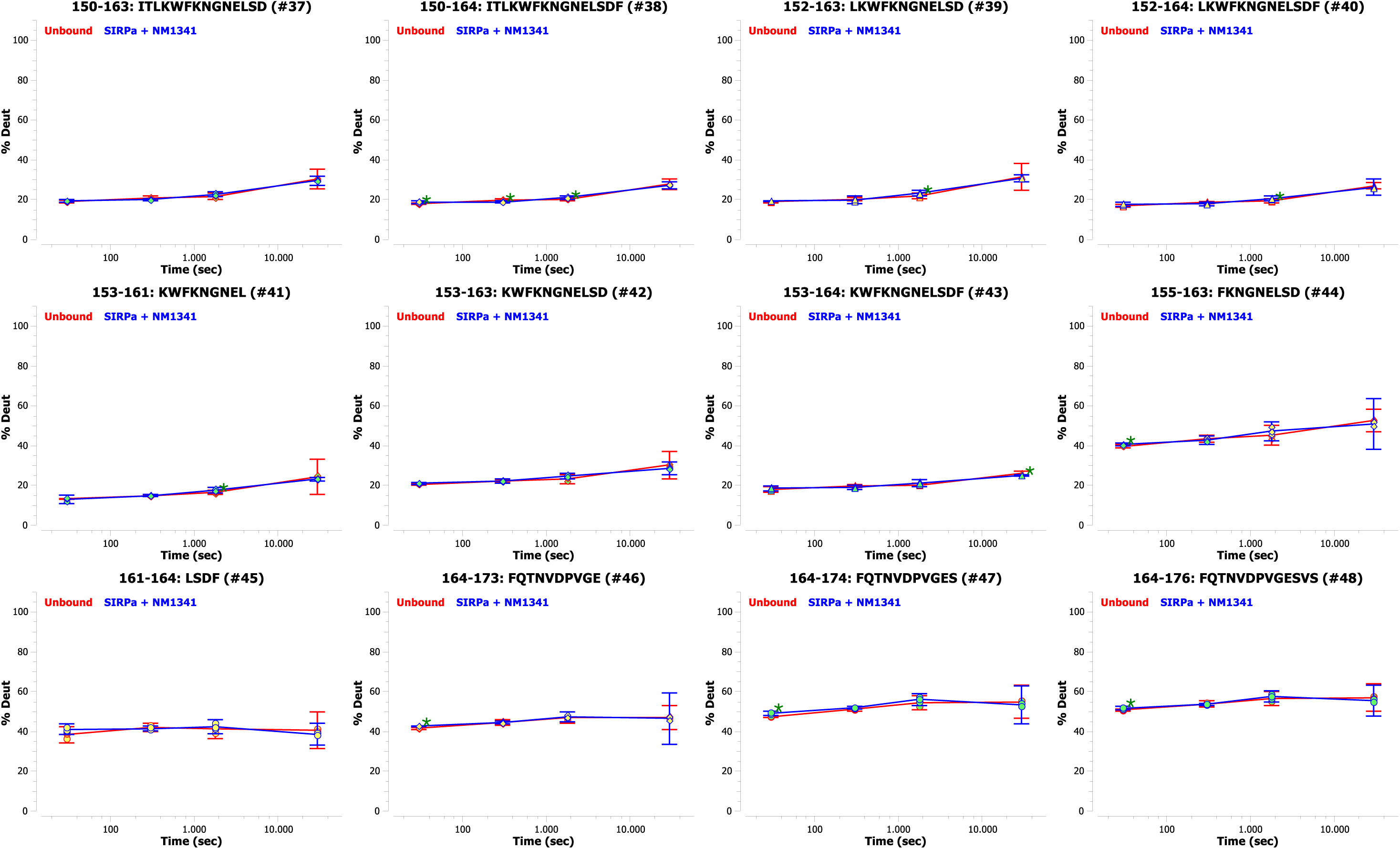
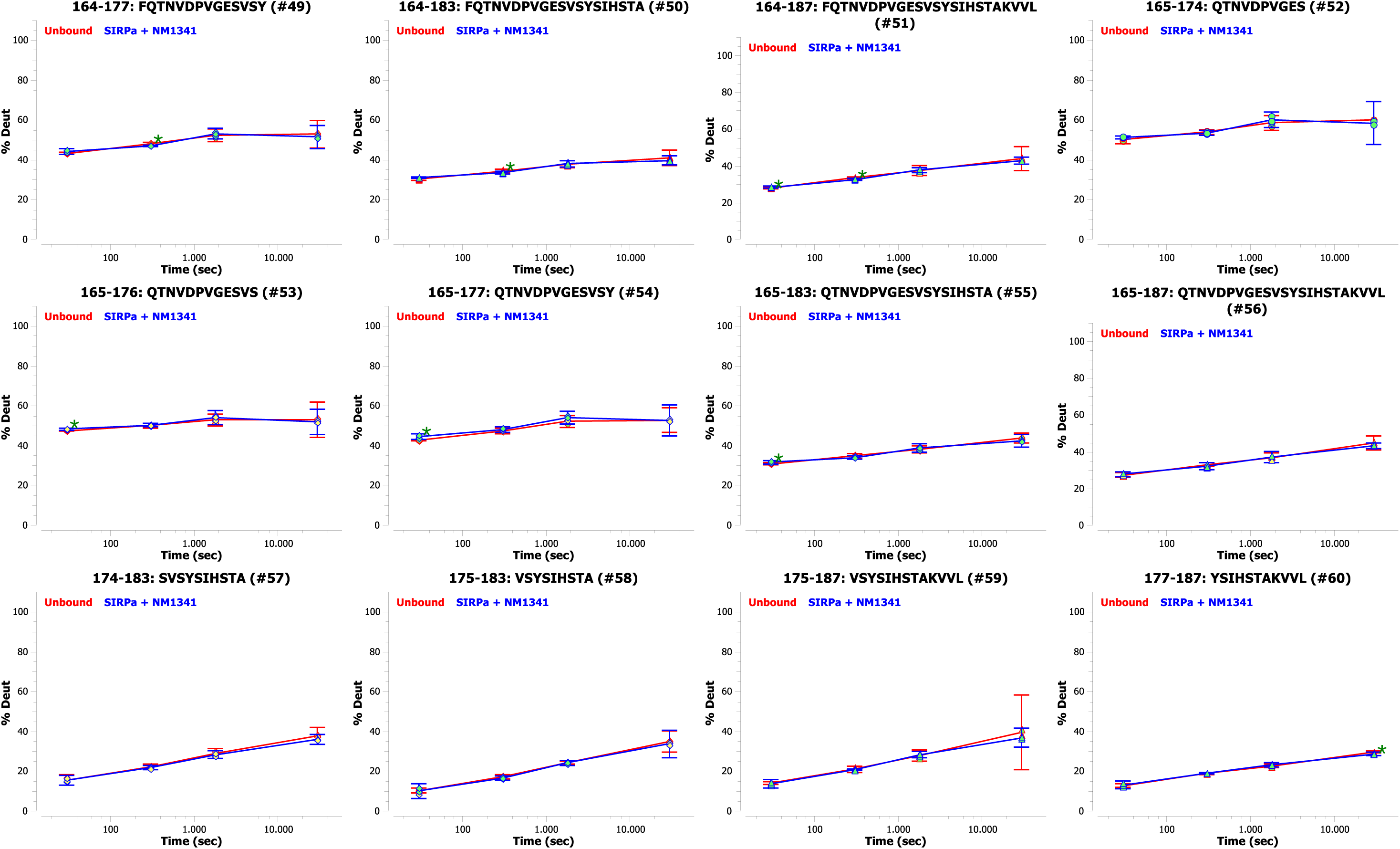
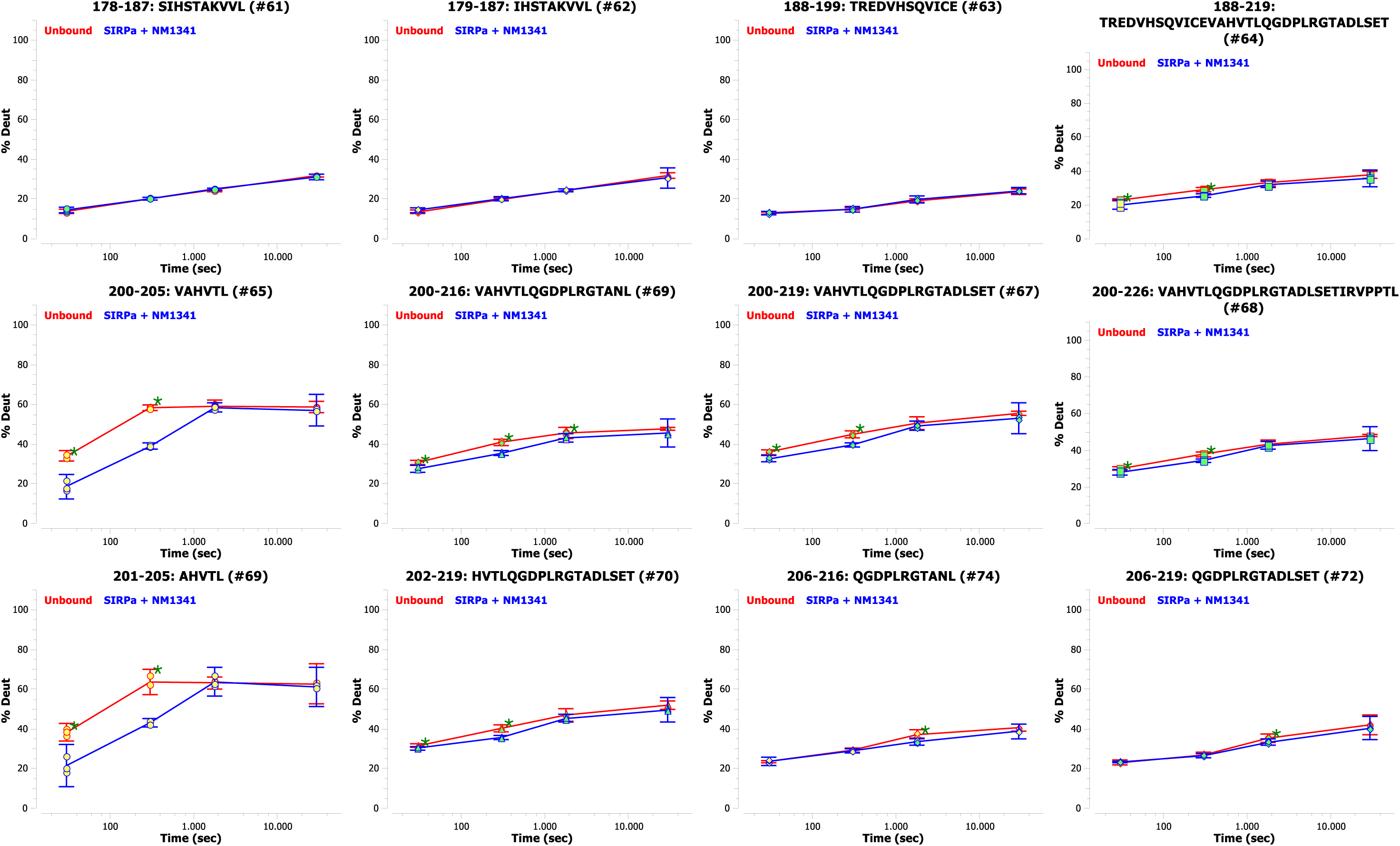
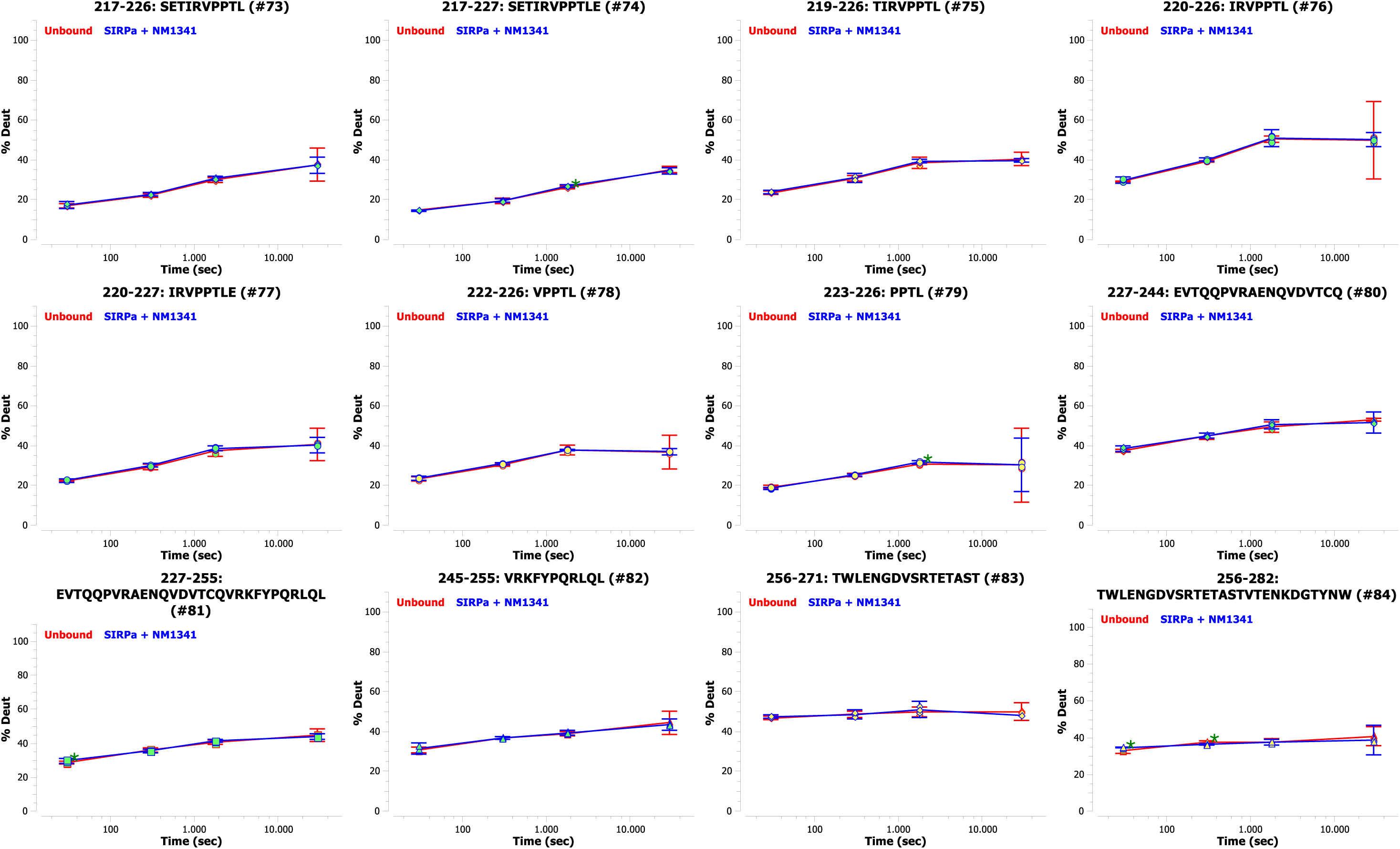
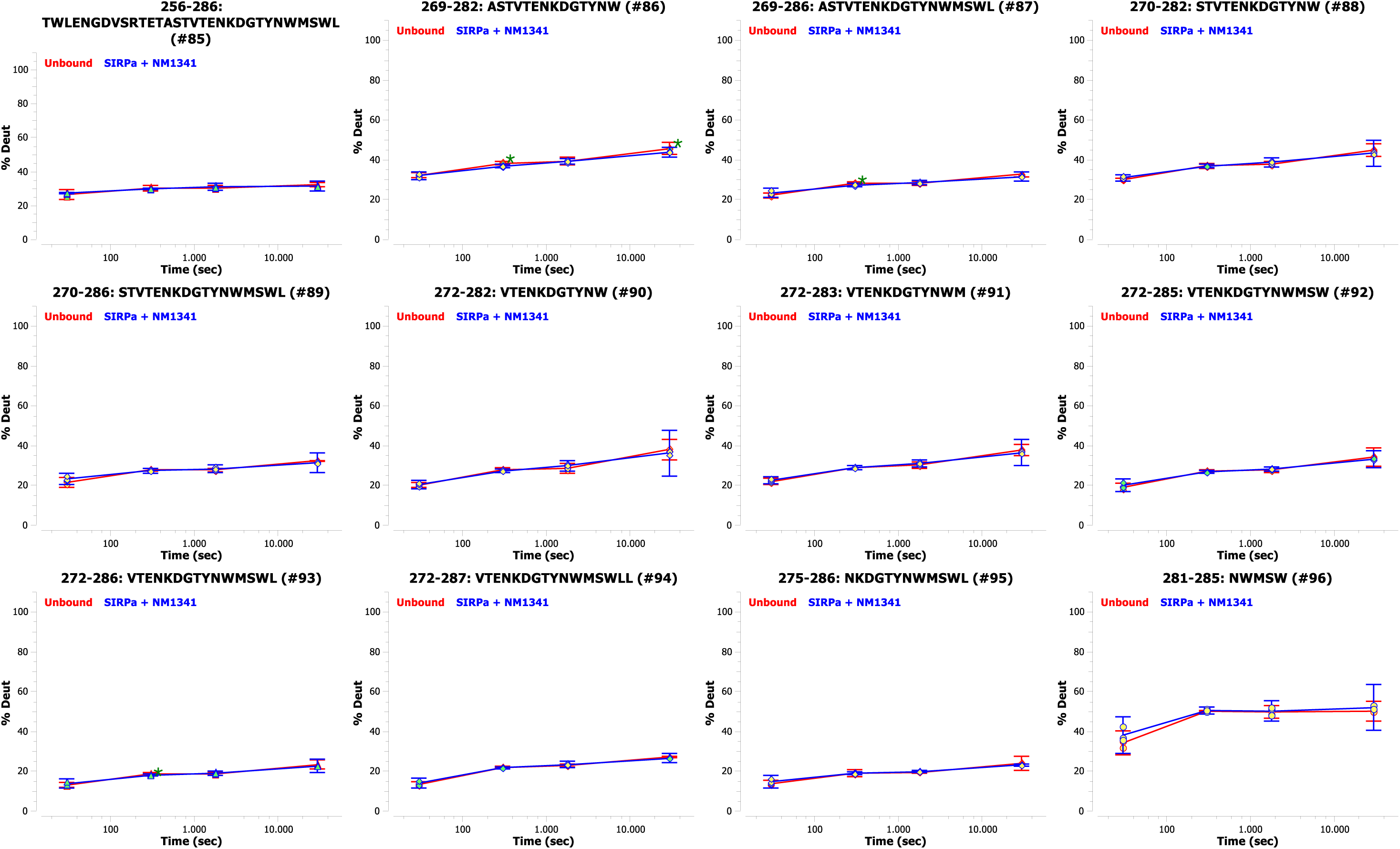
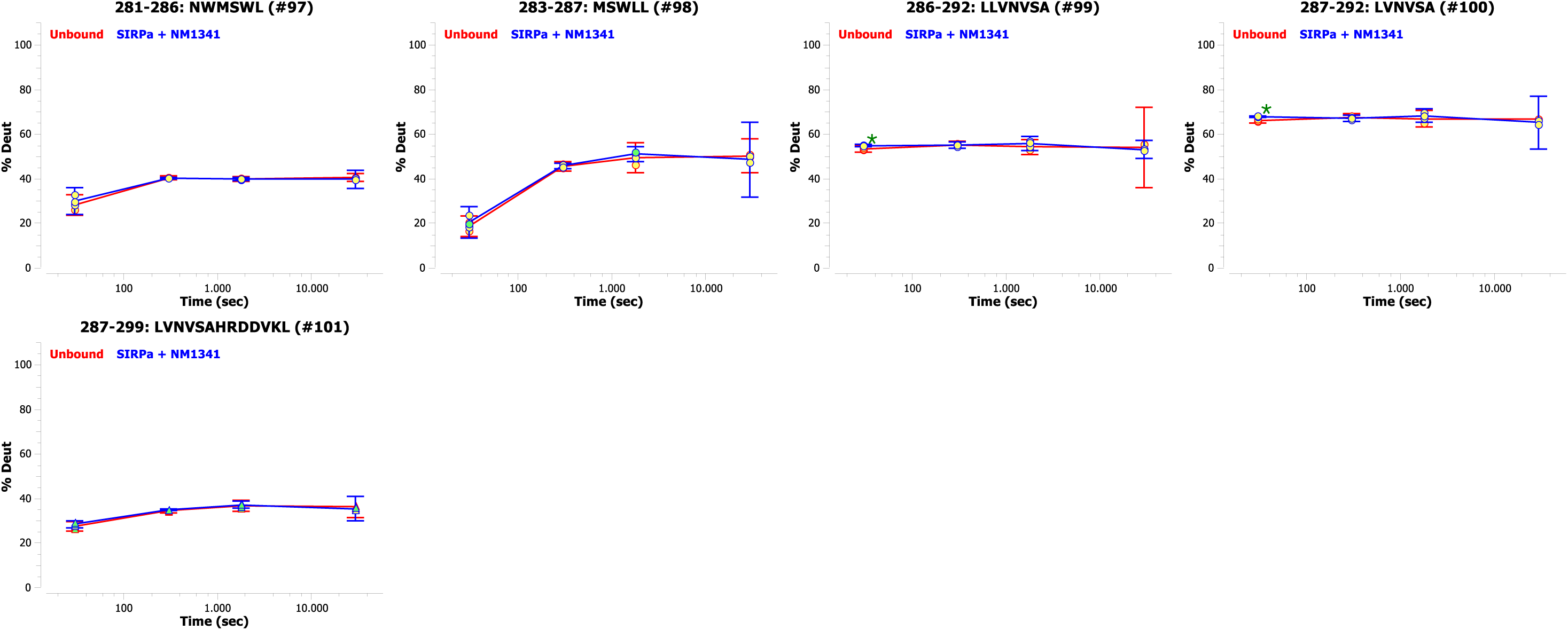
HDX uptake plots of peptic peptides of SIRPα alone and in presence of the nanobody (Identifier NM1341). Deuterium uptake of each peptide is normalized to the exchangeable amino acid residues (number of amino acids minus the first 2 N –terminal residues and proline). Error bars show the significance interval on 95% confidence. Significant difference on basis of the Student’s t-test (p≤0.05) are labeled with an asterisk.

**Figure S7:**
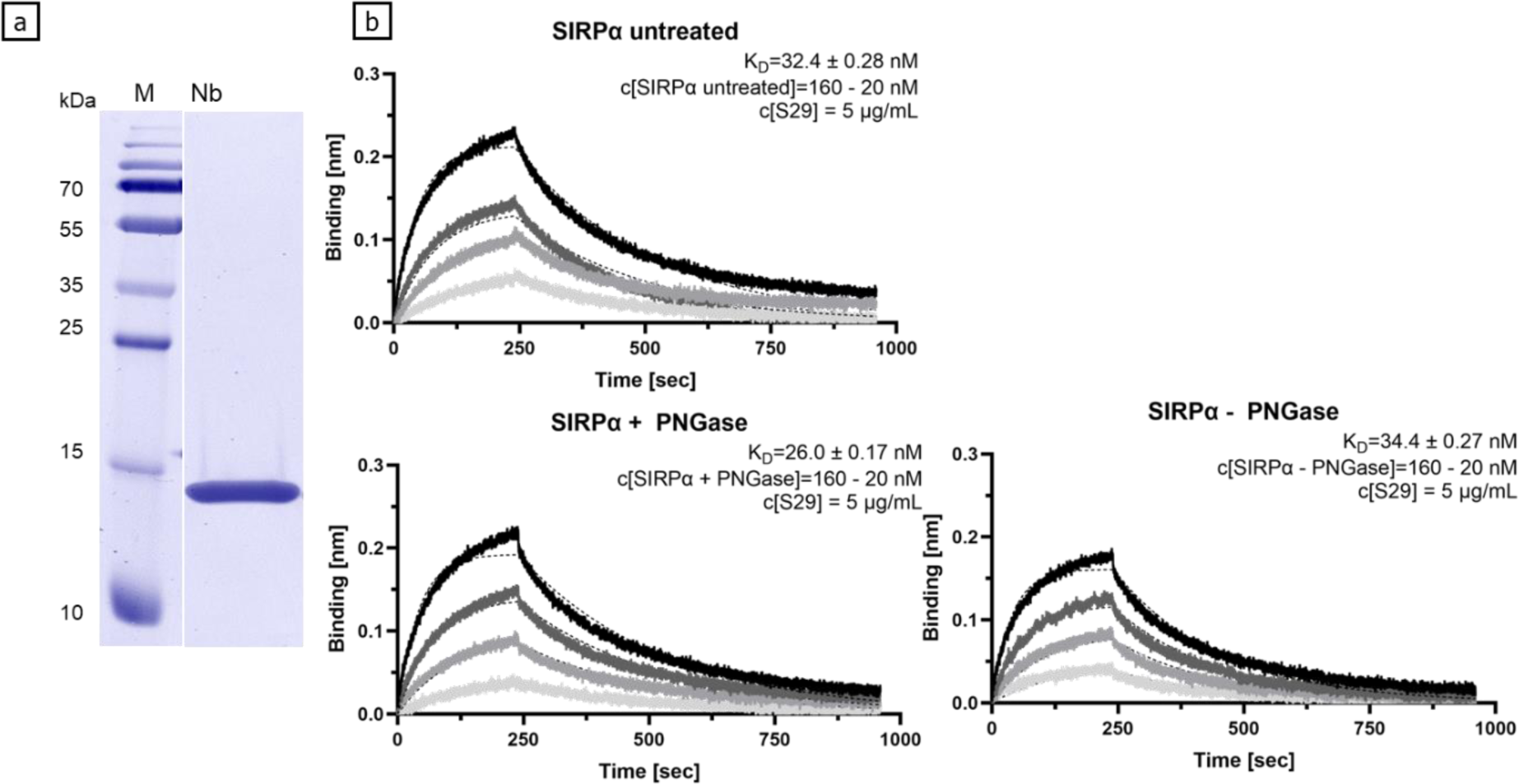
Characterization of the nanobody targeting SIRPα by SDS-PAGE (a), biolayer interferometry (b). (a) SDS-PAGE of purified Nb (M: molecular weight markers) (b) Biolayer interferometry was performed for SIRPα without treatment (upper panel), and with additional deglycosylation overnight at pH 2.5 and 37 °C (lower panel) using PNGase Rc (lower panel, + PNGase). A control was prepared using the same assay conditions without addition of the enzyme (lower panel, -PNGase).

**Figure S8:**
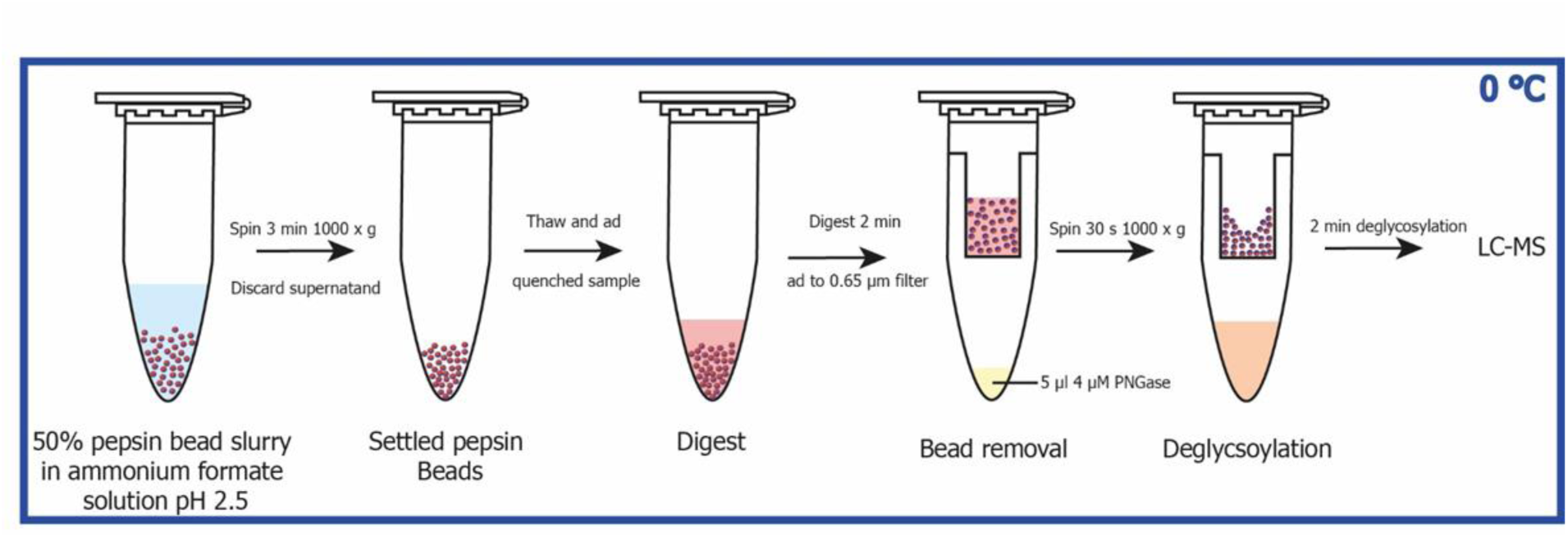
HDX-MS sample preparation with included deglycosylation. The 50% bead slurry was dried by centrifugation and discarding the supernatant. Quenched samples are thawn and incubated with pepsin beads at low temperature (0°C). After 2 min digestion time the bead slurry and solution is transferred to a spin filter in a tube with PNGase RC buffer at low temperature. The peptides from the pepsin digest are quickly spin-filtered to the PNGase solution. After 2 min of PNGase deglycosylation the sample is analyzed by LC-MS. A total of 4.5 min sample preparation time at 0°C minimizes back-exchange of deuterium to hydrogen at the peptide regions of interest. The process was modified from Jensen et al.^33^.

## Supporting Tables

**Table S1:** Masses included in XICs to monitor deglycosylation efficiency using peptide EEQYNSTYR of trastuzumab.

| N-glycan species | Charge | m/z |
| --- | --- | --- |
| A2G0F | 3 | 878.64 - 880.40 |
| A2G1F | 3 | 932.66 - 934.43 |
| A2G2F | 3 | 986.67 - 988.45 |
| A2G0F | 2 | 1317.48 - 1320.09 |
| A2G1F | 2 | 1398.52 - 1401.12 |
| A2G2F | 2 | 1479.53 - 1482.14 |
| Deglycosylated | 2 | 595.70 - 597.31 |
| Deglycosylated | 1 | 1190.45 - 1193.55 |

**Table S2:** Summary of HDX-MS parameters of epitope mapping of the Nb targeting SIRPα as per consensus guidelines^11^.

| Summary of HDX parameters |  |
| --- | --- |
| States | SIRPα & SIRPα bound by the Nb |
| HDX reaction details | 1x PBS pH 7.4, 25 °C, 90% D<sub>2</sub>O |
| Deuteration time points | 0.5, 5, 30 min (n = 3 ) & 500 min (n = 2) |
| Av. back exchange<br>(synthetic peptides; 5 min)<sup>16</sup> | 25% |
| Digestion conditions | 2 min on ice, 30 µl pepsin beads;<br>2 min deglycosylation, 5 µL 4 µM PNGase<br>Rc |
| Average peptide length (AA) / average<br>redundancy (AA) | 14.3 (s<sub>d</sub>=6.7) / 4.1 |
| Number of used peptides/ sequence<br>coverage | 101 / 85.7% |
| Global $\Delta\overline{HX}$ threshold for each time point (at<br>95% confidence) | 0.36 Da |
| Amount of SIRPα:Nb complex during<br>labeling | 99% |

**Table S3:** Sequence homology between the recently published acidic PNGases Dj^37^, Tr^42^ and Rc determined by BLASTp.

|  | Rc | Dj | Tr |
| --- | --- | --- | --- |
| Rc | - | 45% | 47% |
| Dj | 45% | - | 47% |
| Tr | 47% | 47% | - |

## References

(1) Hogg, P. J. Disulfide bonds as switches for protein function. Trends in Biochemical Sciences 2003, 28 (4), 210–214. DOI: 10.1016/s0968-0004(03)00057-4.

(2) Stanley, P.; Schachter, H.; Taniguchi, N. N-Glycans. In Essentials of Glycobiology, 2nd edition, Varki, A., Cummings, R., Esko, J., Freeze, H., Stanley, P., Bertozzi, C., Hart, G., Etzler, M. Eds.; Cold Spring Harbor (NY), 2009.

(3) Sola, R. J.; Griebenow, K. Glycosylation of therapeutic proteins: an effective strategy to optimize efficacy. BioDrugs 2010, 24 (1), 9–21. DOI: 10.2165/11530550-000000000-00000.

(4) Varki, A. Biological roles of glycans. Glycobiology 2017, 27 (1), 3–49. DOI: 10.1093/glycob/cww086.

(5) Reily, C.; Stewart, T. J.; Renfrow, M. B.; Novak, J. Glycosylation in health and disease. Nat. Rev. Nephrol. 2019, 15 (6), 346–366. DOI: 10.1038/s41581-019-0129-4.

(6) Ohtsubo, K.; Marth, J. D. Glycosylation in cellular mechanisms of health and disease. Cell 2006, 126 (5), 855–867. DOI: 10.1016/j.cell.2006.08.019.

(7) Planinc, A.; Bones, J.; Dejaegher, B.; Van Antwerpen, P.; Delporte, C. Glycan characterization of biopharmaceuticals: Updates and perspectives. Anal. Chim. Acta 2016, 921, 13–27. DOI: 10.1016/j.aca.2016.03.049.

(8) Wang, T.; Liu, L.; Voglmeir, J. mAbs N-glycosylation: Implications for biotechnology and analytics. Carbohydr Res 2022, 514, 108541. DOI: 10.1016/j.carres.2022.108541.

(9) Trabjerg, E.; Nazari, Z. E.; Rand, K. D. Conformational analysis of complex protein states by hydrogen/deuterium exchange mass spectrometry (HDX-MS): Challenges and emerging solutions. TrAC Trends in Analytical Chemistry 2018, 106, 125–138. DOI: 10.1016/j.trac.2018.06.008.

(10) Dotz, V.; Haselberg, R.; Shubhakar, A.; Kozak, R. P.; Falck, D.; Rombouts, Y.; Reusch, D.; Somsen, G. W.; Fernandes, D. L.; Wuhrer, M. Mass spectrometry for glycosylation analysis of biopharmaceuticals. TrAC Trends in Analytical Chemistry 2015, 73, 1–9. DOI: 10.1016/j.trac.2015.04.024.

(11) Masson, G. R.; Burke, J. E.; Ahn, N. G.; Anand, G. S.; Borchers, C.; Brier, S.; Bou-Assaf, G. M.; Engen, J. R.; Englander, S. W.; Faber, J.;, et al. Recommendations for performing, interpreting and reporting hydrogen deuterium exchange mass spectrometry (HDX-MS) experiments. Nat. Methods 2019, 16 (7), 595–602. DOI: 10.1038/s41592-019-0459-y.

(12) Merkle, P. S.; Trabjerg, E.; Hongjian, S.; Ferber, M.; Cuendet, M. A.; Jorgensen, T. J. D.; Luescher, I.; Irving, M.; Zoete, V.; Michielin, O.;, et al. Probing the Conformational Dynamics of Affinity-Enhanced T Cell Receptor Variants upon Binding the Peptide-Bound Major Histocompatibility Complex by Hydrogen/Deuterium Exchange Mass Spectrometry. Biochemistry 2021, 60 (11), 859–872. DOI: 10.1021/acs.biochem.1c00035.

(13) Traenkle, B.; Kaiser, P. D.; Pezzana, S.; Richardson, J.; Gramlich, M.; Wagner, T. R.; Seyfried, D.; Weldle, M.; Holz, S.; Parfyonova, Y.;, et al. Single-Domain Antibodies for Targeting, Detection, and In Vivo Imaging of Human CD4+ Cells. Frontiers in Immunology 2021, 12 (5206), Original Research. DOI: 10.3389/fimmu.2021.799910.

(14) Houde, D.; Berkowitz, S. A.; Engen, J. R. The utility of hydrogen/deuterium exchange mass spectrometry in biopharmaceutical comparability studies. J. Pharm. Sci. 2011, 100 (6), 2071–2086. DOI: 10.1002/jps.22432.

(15) Masson, G. R.; Jenkins, M. L.; Burke, J. E. An overview of hydrogen deuterium exchange mass spectrometry (HDX-MS) in drug discovery. Expert Opin. Drug Discov. 2017, 12 (10), 981–994. DOI: 10.1080/17460441.2017.1363734.

(16) Gramlich, M.; Hays, H. C. W.; Crichton, S.; Kaiser, P. D.; Heine, A.; Schneiderhan-Marra, N.; Rothbauer, U.; Stoll, D.; Maier, S.; Zeck, A. HDX-MS for Epitope Characterization of a Therapeutic ANTIBODY Candidate on the Calcium-Binding Protein Annexin-A1. Antibodies 2021, 10 (1). DOI: 10.3390/antib10010011.

(17) Gallagher, E. S.; Hudgens, J. W. Mapping Protein-Ligand Interactions with Proteolytic Fragmentation, Hydrogen/Deuterium Exchange-Mass Spectrometry. Methods Enzymol 2016, 566, 357–404. DOI: 10.1016/bs.mie.2015.08.010.

(18) Chalmers, M. J.; Busby, S. A.; Pascal, B. D.; West, G. M.; Griffin, P. R. Differential hydrogen/deuterium exchange mass spectrometry analysis of protein-ligand interactions. Expert Rev. Proteomics 2011, 8 (1), 43–59. DOI: 10.1586/epr.10.109.

(19) Walters, B. T.; Ricciuti, A.; Mayne, L.; Englander, S. W. Minimizing back exchange in the hydrogen exchange-mass spectrometry experiment. J. Am. Soc. Mass Spectrom. 2012, 23 (12), 2132–2139. DOI: 10.1007/s13361-012-0476-x.

(20) Puchades, C.; Kukrer, B.; Diefenbach, O.; Sneekes-Vriese, E.; Juraszek, J.; Koudstaal, W.; Apetri, A. Epitope mapping of diverse influenza Hemagglutinin drug candidates using HDX-MS. Sci. Rep. 2019, 9 (1), 4735. DOI: 10.1038/s41598-019-41179-0.

(21) Liang, Y.; Guttman, M.; Davenport, T. M.; Hu, S. L.; Lee, K. K. Probing the Impact of Local Structural Dynamics of Conformational Epitopes on Antibody Recognition. Biochemistry 2016, 55 (15), 2197–2213. DOI: 10.1021/acs.biochem.5b01354.

(22) Darula, Z.; Medzihradszky, K. F. Glycan side reaction may compromise ETD-based glycopeptide identification. J. Am. Soc. Mass Spectrom. 2014, 25 (6), 977–987. DOI: 10.1007/s13361-014-0852-9.

(23) Houde, D.; Peng, Y.; Berkowitz, S. A.; Engen, J. R. Post-translational modifications differentially affect IgG1 conformation and receptor binding. Mol. Cell. Proteomics 2010, 9 (8), 1716–1728. DOI: 10.1074/mcp.M900540-MCP200.

(24) Huang, R. Y.; Hudgens, J. W. Effects of desialylation on human alpha1-acid glycoprotein-ligand interactions. Biochemistry 2013, 52 (40), 7127–7136. DOI: 10.1021/bi4011094.

(25) Guttman, M.; Scian, M.; Lee, K. K. Tracking hydrogen/deuterium exchange at glycan sites in glycoproteins by mass spectrometry. Anal. Chem. 2011, 83 (19), 7492–7499. DOI: 10.1021/ac201729v.

(26) Pan, J.; Zhang, S.; Chou, A.; Borchers, C. H. Higher-order structural interrogation of antibodies using middle-down hydrogen/deuterium exchange mass spectrometry. Chem. Sci. 2016, 7 (2), 1480–1486. DOI: 10.1039/c5sc03420e.

(27) Wang, T.; Voglmeir, J. PNGases as valuable tools in glycoprotein analysis. Protein Pept. Lett. 2014, 21 (10), 976–985. DOI: 10.2174/0929866521666140626111237.

(28) Yang, Y.; Wang, G.; Song, T.; Lebrilla, C. B.; Heck, A. J. R. Resolving the micro-heterogeneity and structural integrity of monoclonal antibodies by hybrid mass spectrometric approaches. MAbs 2017, 9 (4), 638–645. DOI: 10.1080/19420862.2017.1290033.

(29) Plummer, T. H.; Elder, J. H.; Alexander, S.; Phelan, A. W.; Tarentino, A. L. Demonstration of peptide:N-glycosidase F activity in endo-beta-N-acetylglucosaminidase F preparations. J. Biol. Chem. 1984, 259 (17), 10700–10704.

(30) Takahashi, N.; Nishibe, H. Some characteristics of a new glycopeptidase acting on aspartylglycosylamine linkages. J. Biochem. 1978, 84 (6), 1467–1473. DOI: 10.1093/oxfordjournals.jbchem.a132270.

(31) Pan, J.; Zhang, S.; Parker, C. E.; Borchers, C. H. Subzero temperature chromatography and top-down mass spectrometry for protein higher-order structure characterization: method validation and application to therapeutic antibodies. J. Am. Chem. Soc. 2014, 136 (37), 13065–13071. DOI: 10.1021/ja507880w.

(32) Wagner, N. D.; Huang, Y.; Liu, T.; Gross, M. L. Post-HDX Deglycosylation of Fc Gamma Receptor IIIa Glycoprotein Enables HDX Characterization of Its Binding Interface with IgG. J. Am. Soc. Mass Spectrom. 2021. DOI: 10.1021/jasms.1c00003.

(33) Jensen, P. F.; Comamala, G.; Trelle, M. B.; Madsen, J. B.; Jorgensen, T. J.; Rand, K. D. Removal of N-Linked Glycosylations at Acidic pH by PNGase A Facilitates Hydrogen/Deuterium Exchange Mass Spectrometry Analysis of N-Linked Glycoproteins. Anal. Chem. 2016, 88 (24), 12479–12488. DOI: 10.1021/acs.analchem.6b03951.

(34) Hamuro, Y.; Coales, S. J. Optimization of Feasibility Stage for Hydrogen/Deuterium Exchange Mass Spectrometry. J. Am. Soc. Mass Spectrom. 2018, 29 (3), 623–629. DOI: 10.1007/s13361-017-1860-3.

(35) Yan, X.; Zhang, H.; Watson, J.; Schimerlik, M. I.; Deinzer, M. L. Hydrogen/deuterium exchange and mass spectrometric analysis of a protein containing multiple disulfide bonds: Solution structure of recombinant macrophage colony stimulating factor-beta (rhM-CSFbeta). Protein Sci 2002, 11 (9), 2113–2124. DOI: 10.1110/ps.0204402.

(36) Zhang, H. M.; McLoughlin, S. M.; Frausto, S. D.; Tang, H.; Emmett, M. R.; Marshall, A. G. Simultaneous reduction and digestion of proteins with disulfide bonds for hydrogen/deuterium exchange monitored by mass spectrometry. Anal Chem 2010, 82 (4), 1450–1454. DOI: 10.1021/ac902550n.

(37) Guo, R. R.; Comamala, G.; Yang, H. H.; Gramlich, M.; Du, Y. M.; Wang, T.; Zeck, A.; Rand, K. D.; Liu, L.; Voglmeir, J. Discovery of Highly Active Recombinant PNGase H(+) Variants Through the Rational Exploration of Unstudied Acidobacterial Genomes. Front. Bioeng. Biotechnol. 2020, 8, 741. DOI: 10.3389/fbioe.2020.00741.

(38) Comamala, G.; Krogh, C. C.; Nielsen, V. S.; Kutter, J. P.; Voglmeir, J.; Rand, K. D. Hydrogen/Deuterium Exchange Mass Spectrometry with Integrated Electrochemical Reduction and Microchip-Enabled Deglycosylation for Epitope Mapping of Heavily Glycosylated and Disulfide-Bonded Proteins. Anal Chem 2021, 93 (49), 16330–16340. DOI: 10.1021/acs.analchem.1c01728.

(39) Weon, H. Y.; Yoo, S. H.; Kim, Y. J.; Lee, C. M.; Kim, B. Y.; Jeon, Y. A.; Hong, S. B.; Anandham, R.; Kwon, S. W. Rudaea cellulosilytica gen. nov., sp. nov., isolated from soil. Int J Syst Evol Microbiol 2009, 59 (Pt 9), 2308–2312. DOI: 10.1099/ijs.0.005165-0.

(40) Sanchez-De Melo, I.; Grassi, P.; Ochoa, F.; Bolivar, J.; Garcia-Cozar, F. J.; Duran-Ruiz, M. C. N-glycosylation profile analysis of Trastuzumab biosimilar candidates by Normal Phase Liquid Chromatography and MALDI-TOF MS approaches. J Proteomics 2015, 127 (Pt B), 225–233. DOI: 10.1016/j.jprot.2015.04.012.

(41) Wang, T.; Zheng, S. L.; Liu, L.; Voglmeir, J. Development of a colorimetric PNGase activity assay. Carbohydr Res 2019, 472, 58–64. DOI: 10.1016/j.carres.2018.11.007.

(42) Wang, T.; Cai, Z. P.; Gu, X. Q.; Ma, H. Y.; Du, Y. M.; Huang, K.; Voglmeir, J.; Liu, L. Discovery and characterization of a novel extremely acidic bacterial N-glycanase with combined advantages of PNGase F and A. Biosci. Rep. 2014, 34 (6). DOI: 10.1042/BSR20140148.

(43) Joao, H. C.; Dwek, R. A. Effects of glycosylation on protein structure and dynamics in ribonuclease B and some of its individual glycoforms. Eur J Biochem 1993, 218 (1), 239–244. DOI: 10.1111/j.1432-1033.1993.tb18370.x.

(44) Windwarder, M.; Altmann, F. Site-specific analysis of the O-glycosylation of bovine fetuin by electron-transfer dissociation mass spectrometry. J Proteomics 2014, 108, 258–268. DOI: 10.1016/j.jprot.2014.05.022.

(45) Hatherley, D.; Graham, S. C.; Harlos, K.; Stuart, D. I.; Barclay, A. N. Structure of signal-regulatory protein alpha: a link to antigen receptor evolution. J. Biol. Chem. 2009, 284 (39), 26613–22619. DOI: 10.1074/jbc.M109.017566.

(46) Oliveira, T.; Thaysen-Andersen, M.; Packer, N. H.; Kolarich, D. The Hitchhiker’s guide to glycoproteomics. Biochem Soc Trans 2021, 49 (4), 1643–1662. DOI: 10.1042/BST20200879.

(47) Altmann, F.; Schweiszer, S.; Weber, C. Kinetic comparison of peptide: N-glycosidases F and A reveals several differences in substrate specificity. Glycoconj. J. 1995, 12 (1), 84–93. DOI: 10.1007/BF00731873.

(48) Ruiz-May, E.; Thannhauser, T. W.; Zhang, S.; Rose, J. K. Analytical technologies for identification and characterization of the plant N-glycoproteome. Front Plant Sci 2012, 3, 150. DOI: 10.3389/fpls.2012.00150.

(49) Staudacher, E. Fucose in N-glycans: from plant to man. Biochimica et Biophysica Acta BBA) 1999, 1473 (1), 216–236. DOI: 10.1016/s0304-4165(99)00181-6.

(50) Wilson, I. B.; Zeleny, R.; Kolarich, D.; Staudacher, E.; Stroop, C. J.; Kamerling, J. P.; Altmann, F. Analysis of Asn-linked glycans from vegetable foodstuffs: widespread occurrence of Lewis a, core alpha1,3-linked fucose and xylose substitutions. Glycobiology 2001, 11 (4), 261–274. DOI: 10.1093/glycob/11.4.261.

(51) Zeck, A.; Regula, J. T.; Larraillet, V.; Mautz, B.; Popp, O.; Gopfert, U.; Wiegeshoff, F.; Vollertsen, U. E.; Gorr, I. H.; Koll, H.;, et al. Low level sequence variant analysis of recombinant proteins: an optimized approach. PLoS One 2012, 7 (7), e40328. DOI: 10.1371/journal.pone.0040328.

(52) Wagner, T. R.; Ostertag, E.; Kaiser, P. D.; Gramlich, M.; Ruetalo, N.; Junker, D.; Haering, J.; Traenkle, B.; Becker, M.; Dulovic, A.;, et al. NeutrobodyPlex-monitoring SARS-CoV-2 neutralizing immune responses using nanobodies. EMBO Rep. 2021, 22 (5). DOI: 10.15252/embr.202052325.

(53) Kochert, B. A.; Iacob, R. E.; Wales, T. E.; Makriyannis, A.; Engen, J. R. Hydrogen-Deuterium Exchange Mass Spectrometry to Study Protein Complexes. Methods Mol. Biol. 2018, 1764, 153–171. DOI: 10.1007/978-1-4939-7759-8_10.

